# Hematopoietic stem cell gene therapy improves outcomes in a clinically relevant mouse model of Multiple Sulfatase Deficiency

**DOI:** 10.1101/2024.03.08.584099

**Authors:** Vi Pham, Lucas Tricoli, Xinying Hong, Parith Wongkittichote, Carlo Castruccio Castracani, Amaliris Guerra, Lars Schlotawa, Laura A. Adang, Amanda Kuhs, Margaret M. Cassidy, Owen Kane, Emily Tsai, Maximiliano Presa, Cathleen Lutz, Stefano B. Rivella, Rebecca C. Ahrens-Nicklas

## Abstract

Multiple sulfatase deficiency (MSD) is a severe, lysosomal storage disorder caused by pathogenic variants in the gene *SUMF1,* encoding the sulfatase modifying factor formylglycine-generating enzyme. Patients with MSD exhibit functional deficiencies in all cellular sulfatases. The inability of sulfatases to break down their substrates leads to progressive and multi-systemic complications in patients, similar to those seen in single-sulfatase disorders such as metachromatic leukodystrophy and mucopolysaccharidoses IIIA. Here, we aimed to determine if hematopoietic stem cell transplant with *ex vivo SUMF1* lentiviral gene therapy could improve outcomes in a clinically relevant mouse model of MSD. We first tested our approach in MSD patient-derived cells and found that our *SUMF1* lentiviral vector improved protein expression, sulfatase activities, and glycosaminoglycan accumulation. *In vivo*, we found that our gene therapy approach rescued biochemical deficits, including sulfatase activity and glycosaminoglycan accumulation, in affected organs of MSD mice treated post-symptom onset. In addition, treated mice demonstrated improved neuroinflammation and neurocognitive function. Together, these findings suggest that *SUMF1* HSCT-GT can improve both biochemical and functional disease markers in the MSD mouse.

## INTRODUCTION

Multiple sulfatase deficiency (MSD) is an ultra-rare, inherited lysosomal storage disorder (LSD) characterized by the functional deficiency of all sulfatase enzymes. The estimated incidence of MSD is 1 in 500,000 individuals.^1^ MSD arises from pathogenic variants in *SUMF1,*^2,3^ the gene encoding formylglycine-generating enzyme (FGE).^4,5^ FGE is the rate-limiting modifying factor required for activation of all sulfatases in humans.^6^ Without this crucial enzyme, newly synthesized sulfatases are not activated after translation in the endoplasmic reticulum.^7–9^ Loss of sulfatase activities induces the accumulation of various toxic sulfated substrates throughout the body, including sulfatides and glycosaminoglycans (GAGs). Since the discovery of the *SUMF1* gene, at least 21 nonsense mutations and 32 missense mutations over the entire protein have been published.^10^ Most MSD cases are due to hypomorphic variants in *SUMF1* that result in varying degrees of residual sulfatase activities, as cases of null biallelic variants are thought to be incompatible with survival past the newborn period.^11^ Symptoms in patients arise from the additive effects of each individual sulfatase deficiency, which include the sulfatases responsible for seven types of LSDs (i.e. metachromatic leukodystrophy (MLD) and several mucopolysaccharidoses (MPS) subtypes).^12^ Patients with MSD suffer from neurologic regression and progressive somatic symptoms, including respiratory complications, hepatomegaly, bone abnormalities, cardiac dysfunction and hearing loss (for a full clinical review see^13^). The median age of death is approximately 13 years.^14^ As there are no approved disease-modifying therapies for MSD, there is a significant unmet need for new treatments.

Through ongoing MSD natural history studies, we have found that neuromuscular complications are a prominent feature of MSD, and neurological symptoms are a major cause of morbidity. Patients experience severe psychomotor regression and neurological manifestations including developmental delay, seizures, and psychiatric abnormalities.^14^ In addition, neuroinflammation is a characteristic feature of LSDs with central nervous system (CNS) involvement, such as MSD, and may contribute to the pathology.^15^ MSD strongly resembles the neuronopathic, monogenic sulfatase disorders including MLD, Sanfilippo syndrome (MPS IIIA and D), and MPS II, both biochemically and clinically.^14,16^ Hematopoietic stem cell transplantation (HSCT) has been trialed in these disorders with varying success. The rationale of using HSCT to treat neurological manifestations is that donor bone-marrow derived microglia may populate the brain, attenuate inflammation, and secrete activated sulfatases to neighboring cells through cross-correction.^17^ While this approach shows benefit in early-stage MPS I and pre-symptomatic juvenile MLD patients,^18–22^ HSCT alone is insufficient to prevent neurologic progression in infantile MLD and several other MPS disorders.^23^

An alternative approach combining HSCT with gene therapy (HSCT-GT) has proven to be more beneficial and is currently in development for several LSDs of the brain. *Ex vivo* lentiviral gene therapy with hematopoietic stem cell transplant is approved by the European Medicines Agency for MLD^24,25^ and is currently in clinical trials for MPS IIIA (NCT04201405).^26,27^ This approach is also being developed for other related MPS disorders.^28–30^ It has been hypothesized that HSCT-GT may be superior to HSCT alone, likely because the supraphysiologic enzyme levels provided through gene therapy lead to more robust cross-correction.^17^ Notably, transplant-related complications are reduced in HSCT-GT compared to standard allogeneic HSCT, as the patient receives their own cells (i.e. an autologous transplant).

Given the mechanistic and phenotypic overlap of MSD with MLD and MPS IIIA, we hypothesize that HSCT-based *ex vivo* gene therapy may provide therapeutic benefit in MSD. In this initial proof-of-concept study, we demonstrate that *SUMF1* HSCT-GT can reduce storage accumulation throughout the CNS and periphery, attenuate neuroinflammation, and improve behavioral markers of CNS disease in MSD mice.

## RESULTS

### *SUMF1* lentiviral vector improves biochemical and cellular MSD phenotypes *in vitro*

To develop a therapeutic approach that could provide clinical benefits to many MSD patients and address the wide range of pathogenic *SUMF1* variants, we designed a lentiviral vector for HSCT-GT that expresses a full, functional copy of the human *SUMF1* gene (**Figure 1A**). To drive strong and ubiquitous expression of *SUMF1*, our lentiviral vector includes the human elongation factor 1-alpha (EF-1a) promoter, which is known to achieve robust, constitutive, and durable expression in cells and tissues.^31^ We also included ankyrin and foamy viral insulators to enhance safety and reduce the position effects of transcriptional silencing.^32^ Finally, our construct contains a woodchuck hepatitis virus posttranscriptional regulatory element (WPRE)^33^ downstream of the 3’ LTR, followed by a polyadenylation sequence to preserve viral genome integrity.^34^ This vector backbone is further characterized in the companion paper submitted by Tricoli et al.

**Figure 1.**
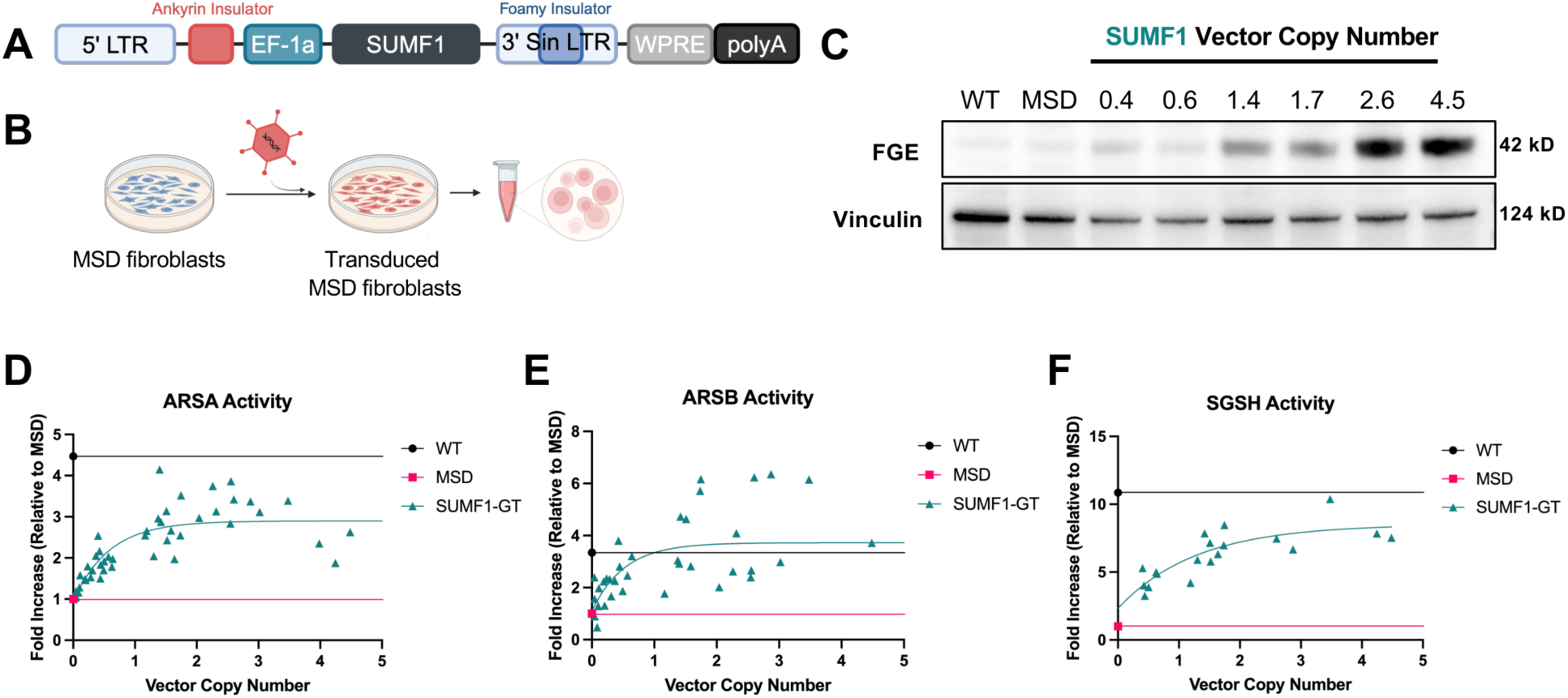
*SUMF1* lentiviral vector improves FGE protein expression and sulfatase activities *in vitro*. A) Lentiviral vector construct encoding human *SUMF1* gene, driven by the human elongation factor 1 alpha (EF-1a) promoter and containing ankyrin and foamy insulators. 5’ LTR: 5’ long terminal repeat; 3’ Sin LTR: 3’ self-inactivating long terminal repeat; WPRE: woodchuck hepatitis virus post-transcriptional regulatory element; polyA: polyadenylation sequence. B) Workflow of *in vitro* experiments. Immortalized MSD patient fibroblasts were transduced with our *SUMF1* lentiviral vector before being harvested for downstream analyses. C) Western blot analysis of FGE protein expression in wild-type (WT) cells, non-transduced MSD cells (MSD), and MSD cells transduced with the *SUMF1* lentiviral vector over a range of vector copy numbers (0.4-4.5) shows increasing protein expression with increasing integrations. Vinculin was used as a loading control. D) ARSA activity (reported as fold-increase relative to non-transduced MSD cells) of WT cells, non-transduced MSD cells (MSD), and MSD cells transduced with *SUMF1* lentiviral vector over a range of vector copy numbers (SUMF1-GT). E) ARSB activity (reported as fold-increase relative to non-transduced MSD cells) of WT cells, non-transduced MSD cells (MSD), and MSD cells transduced with *SUMF1* lentiviral vector over a range of vector copy numbers (SUMF1-GT). F) SGSH activity (reported as fold-increase relative to non-transduced MSD cells) of WT cells, non-transduced MSD cells (MSD), and MSD cells transduced with *SUMF1* lentiviral vector over a range of vector copy numbers (SUMF1-GT). For all sulfatase activity panels, WT and non-transduced MSD values represent the mean of 10 biological replicates for each group. SUMF1-GT individual biological replicate values are shown. For all sulfatase activity panels, the data are fit to an exponential regression line.

We first characterized our *SUMF1*-expressing lentiviral vector *in vitro* by transducing an immortalized, MSD patient-derived primary fibroblast line that harbors a homozygous missense mutation in *SUMF1* (c.463C>T, p.Ser155Pro; **Figure 1B**).^3,35^ We transduced the cells with varying amounts of lentiviral vector to achieve a range of vector copy number (VCN) integrations and examine dose-dependent effects. To validate the expression of our *SUMF1* transgene, we first measured FGE protein expression by Western blot (**Figure 1C**). Both wild-type (WT) and untreated MSD (MSD) cells did not express high levels of endogenous FGE. However, transduction with our *SUMF1* lentiviral vector dramatically increased FGE expression in MSD cells, correlating with increasing lentiviral vector integrations (**Figure 1C**).

Next, we assessed the functional effects of our *SUMF1* lentiviral vector on the biochemistry of MSD patient-derived cells. Post-translational modification by FGE is a key step needed to activate every sulfatase in the cell. Therefore, pathogenic variants in *SUMF1* leading to degradation-prone forms of FGE result in the functional deficiency of all sulfatases in MSD patients.^9,16^ To determine if overexpressing functional FGE with our *SUMF1* lentiviral vector rescues the activities of downstream sulfatases in transduced MSD cells, we measured the activities of three sulfatases: arylsulfatase A (ARSA), arylsulfatase B (ARSB), and sulfamidase (SGSH). Representative of biochemical MSD phenotypes, untreated MSD fibroblasts (MSD) demonstrate 4.5-fold, 3-fold, and 11-fold less activity than WT fibroblasts for ARSA, ARSB, and SGSH, respectively (**Figures 1D-F**). Transduction with our *SUMF1* gene therapy lentiviral vector (SUMF1-GT) increased ARSA activity in MSD fibroblasts, with the highest activity reaching about 4-fold compared to untreated cells. Our *SUMF1* lentiviral vector also enhanced ARSB and SGSH activity in MSD fibroblasts to about 6-fold and 10-fold relative to untreated cells, respectively. Interestingly, we observed a plateau in sulfatase activities, perhaps due to the rate-limiting abundance of target sulfatases. Importantly though, we observed a VCN-dependent increase in sulfatase activities over the clinically relevant range of VCNs (0.5-2).

Finally, we examined the ability of our *SUMF1* lentiviral vector to rescue pathological markers of disease. A hallmark feature of MSD and related single-sulfatase disorders is the accumulation of GAGs in patient cells and tissues due to the inability of sulfatases to break them down.^36^ We detected six different GAG subspecies in MSD cells that all showed significant accumulation compared to WT cells by mass spectrometry using the endogenous, non-reducing end (NRE) biomarker method (**Figures 2A-D**).^36,37^ These GAG subspecies are associated with MPS I/II, IIIA, IIID, and VI, single-sulfatase disorders closely related to MSD. To investigate if our *SUMF1* lentiviral vector could rescue this GAG accumulation, we quantified the levels of GAG subspecies in MSD cells after transduction. We found that MSD cells transduced with our lentiviral vector exhibited significantly decreased levels of GAG subspecies associated with MPS I/II (UA-HNAc-UA-1S), MPS IIIA (HN-UA-1S, HN-UA-HNAc-UA-1S, and HN-UA-HNAc-UA-2S), and MPS IIID (GlcNAc-6S) down to WT levels. These data indicate that our *SUMF1* lentiviral vector can rescue the accumulation of multiple GAG subspecies in MSD patient cells. Overall, these findings demonstrate the ability of our *SUMF1* lentiviral vector to increase FGE protein expression, increase sulfatase activities, and reduce GAG accumulation in MSD cells.

**Figure 2.**
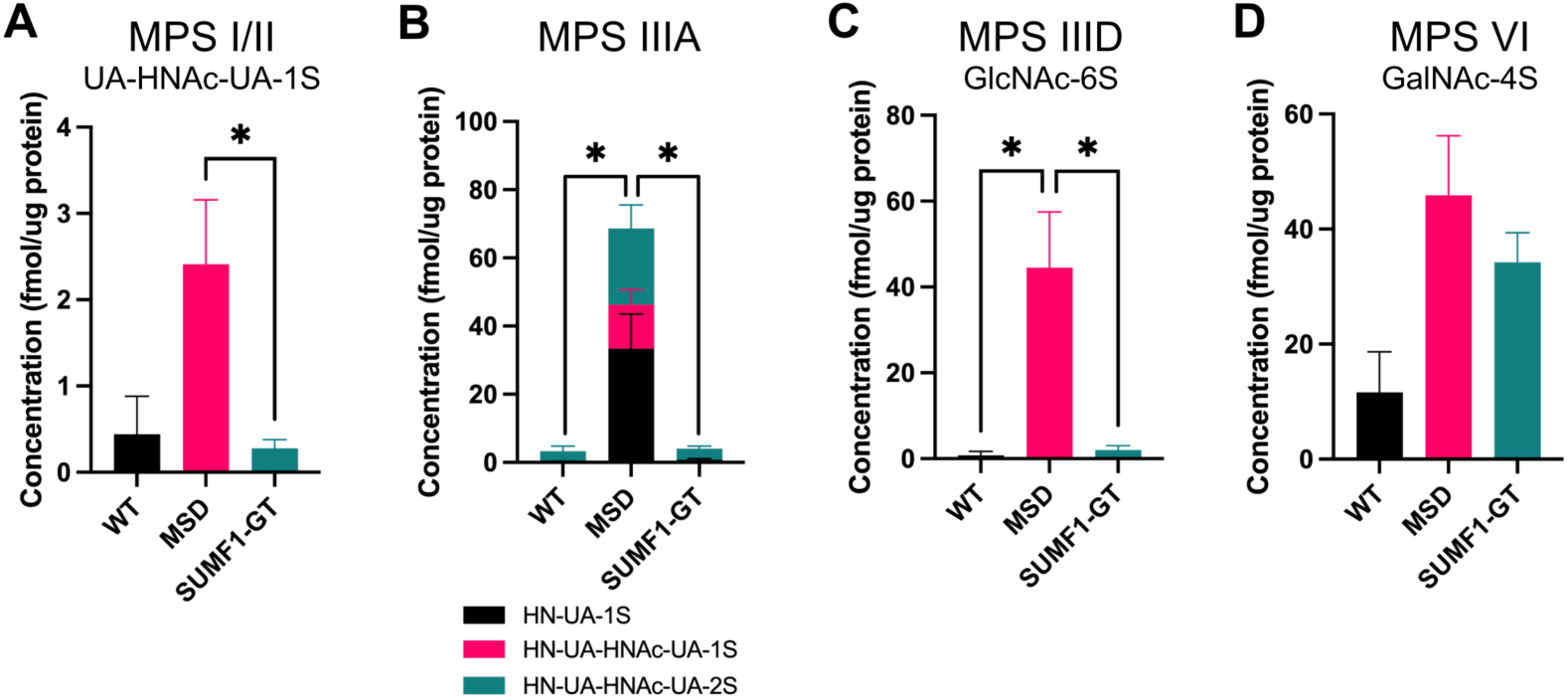
*SUMF1* lentiviral vector rescues GAG accumulation *in vitro*. A) Quantification of GAG subspecies, UA-HNAc-UA-1S, associated with MPS I/II. B) Quantification of GAG subspecies associated with MPS IIIA: HN-UA-1S, HN-UA-HNAc-UA-1S, and HN-UA-HNAc-UA-2S. C) Quantification of GAG subspecies associated with MPS IIID, GlcNAc-6S. D) Quantification of GAG subspecies associated with MPS VI, GalNAc-4S. For all panels, the total concentration of GAGs was normalized to μg of total protein from cell lysates. Total protein was measured by BCA assay. *N* = 2-5 biological replicates, one-way ANOVA, followed by Bonferroni’s multiple comparisons test to compare each group to the untreated MSD control group. For all panels: *p < 0.05. Data are shown as mean ± SEM.

### Hematopoietic stem cell transplant with *SUMF1* lentiviral gene therapy is safe and durable

To determine the safety and efficacy of our *SUMF1* lentiviral vector *in vivo*, we next performed HSCT-GT in both WT mice and a mouse model of MSD. The recently-developed mouse model of MSD harbors a clinically relevant pathogenic variant in the mouse *Sumf1* gene (p.Ser153Pro, corresponding to the human *SUMF1* variant p.Ser155Pro).^38^ Because the original *Sumf1-/-* knockout mouse model of MSD exhibits severe biochemical phenotypes and neonatal lethality, that model is not appropriate to conduct our *ex vivo* gene therapy studies as it does not accurately recapitulate the human disease, nor do mice survive long enough to perform bone marrow transplants that would parallel human therapeutic approaches.^39^

To perform HSCT-GT in the mouse, lineage-negative hematopoietic stem cells (HSCs) were first extracted and isolated from donor mice (**Figure 3A**). Lineage-negative HSCs were then transduced *ex vivo* with our *SUMF1* lentiviral vector before being transplanted into irradiated, 8-week-old recipient mice. Mice of the opposite sex were used as donors/recipients to measure engraftment efficiency by ddPCR for the male-specific gene *Zfy1*. Engraftment was measured in blood at 2 months post-transplant and in the bone marrow at 4 months post-transplant (study endpoint). All transplanted mice reached engraftment levels of 88-100% donor cells by 4 months post-transplant, demonstrating the engraftment efficacy of our HSCT-GT approach in MSD mice (**Figure 3B**).

**Figure 3.**
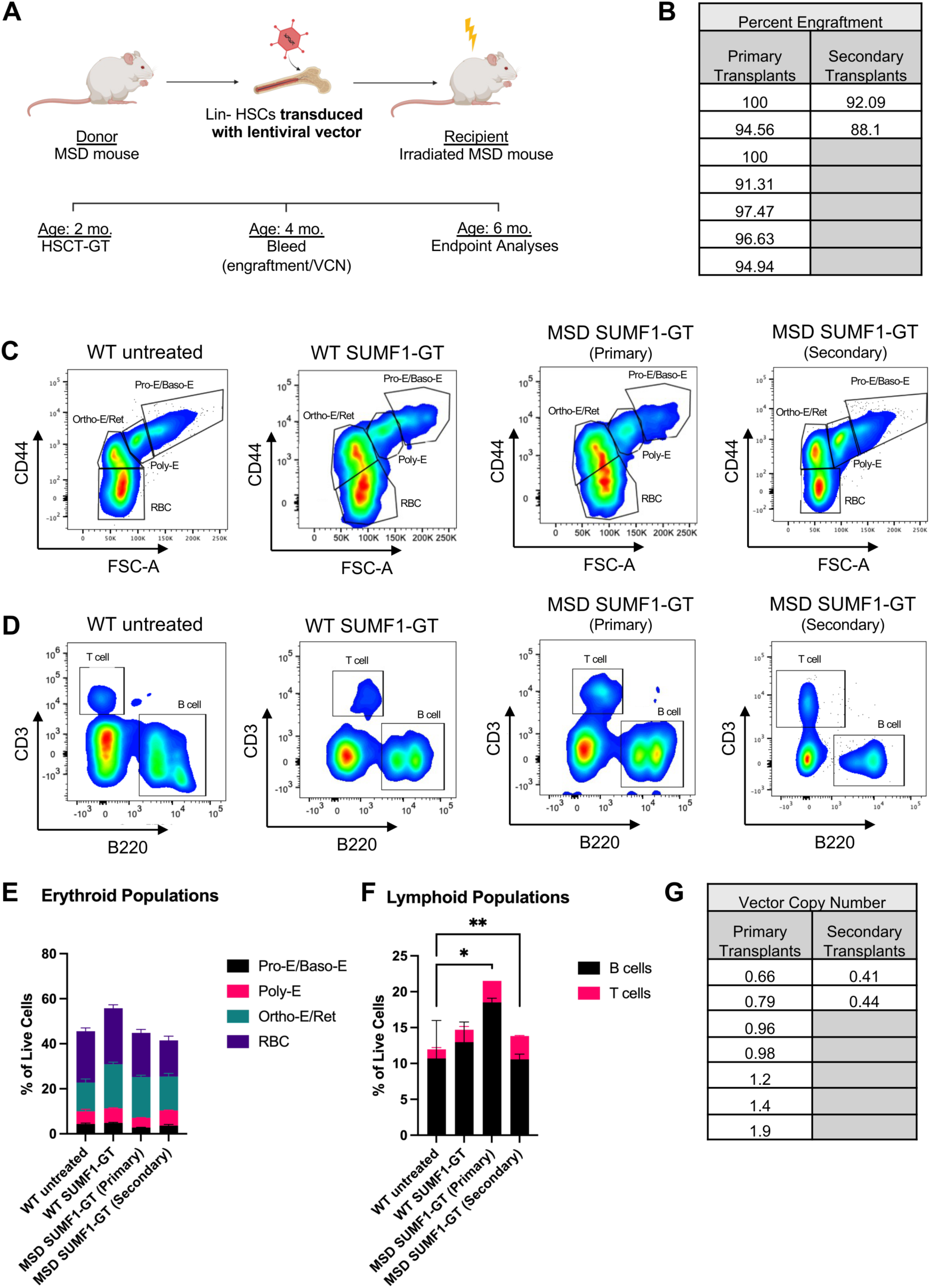
Hematopoietic stem cell transplant with *SUMF1* gene therapy is stable and does not affect hematopoiesis *in vivo*. A) Workflow and timeline of *in vivo* HSCT-GT experiments. Mice receive HSCT-GT at 8 weeks of age, engraftment and VCN are measured at 2 months post-transplant by retro-orbital bleed, and endpoint analyses are conducted 4 months post-transplant. B) Percentages of donor cell engraftment for *SUMF1* HSCT-GT primary and secondary mice. C) Representative flow panels of erythroid populations (CD44+) for untreated WT mice (WT untreated), WT mice receiving *SUMF1* HSCT-GT (WT SUMF1-GT), MSD mice receiving primary *SUMF1* HSCT-GT (MSD SUMF1-GT (Primary)), and MSD mice receiving secondary *SUMF1* HSCT-GT (MSD SUMF1-GT (Secondary)). D) Representative flow panels of lymphoid populations (B220+ or CD3+) for untreated WT mice (WT untreated), WT mice receiving *SUMF1* HSCT-GT (WT SUMF1-GT), MSD mice receiving primary *SUMF1* HSCT-GT (MSD SUMF1-GT (Primary)), and MSD mice receiving secondary *SUMF1* HSCT-GT (MSD SUMF1-GT (Secondary)). E) Quantification of erythroid populations from flow analyses, displayed as percentage of the total live cell population. *N* = 2-5 mice, one-way ANOVA for each cell population followed by Bonferroni’s multiple comparisons test to compare each group to the WT untreated group. Data are represented as mean ± SEM. No significant differences between groups. F) Quantification of lymphoid populations from flow analyses, displayed as percentage of the total live cell population. *N* = 2-5 mice, one-way ANOVA for each cell population followed by Bonferroni’s multiple comparisons test to compare each group to the WT untreated group. No significant differences in B cell populations between groups. T cell populations in MSD SUMF1-GT (Primary) and MSD SUMF1-GT (Secondary) mice were greater than WT untreated mice. *p < 0.05, **p <0.01. Data are represented as mean ± SEM. G) Vector copy numbers of *SUMF1* HSCT-GT primary and secondary mice.

We first established the safety of our *SUMF1* lentiviral vector itself by conducting *SUMF1* HSCT-GT in a WT background (WT SUMF1-GT). In this group, WT donor HSCs were transduced with our *SUMF1* lentiviral vector before being transplanted into WT recipient mice. We then examined both safety and efficacy of our *SUMF1* lentiviral vector in the MSD background by conducting *SUMF1* HSCT-GT in MSD donor and recipient mice (MSD SUMF1-GT (Primary)). Finally, to determine the durability of our approach, we conducted secondary transplants by extracting bone marrow from 6-month-old MSD mice that received *SUMF1* gene therapy and transplanting the already-transduced bone marrow cells into naïve, irradiated MSD mice at 8-weeks of age (MSD SUMF1-GT (Secondary)). All animals survived in good condition to study endpoint and demonstrated no outward signs of vector-related toxicity.

For all groups, we assessed the ability of *SUMF1-*transduced donor HSCs to differentiate into erythroid (**Figures 3C and 3E**) and lymphoid (**Figures 3D and 3F**) populations by conducting flow cytometry analyses on bone marrow at study endpoint (4 months post-transplant). HSCs from mice of all groups were successfully able to differentiate into erythroid populations, including pro-erythroblasts and basophilic erythroblasts (Pro-E/Baso-E), polychromatic erythroblasts (Poly-E), orthochromatic erythroblasts and (Ortho-E/Ret), and red blood cells (RBCs), in comparable proportions to untreated WT mice (**Figure 3E**). For lymphoid populations, there were no statistically significant differences in B cell populations of treated groups as compared to untreated WT mice (**Figure 3F**). There was a slight increase in T cell populations in MSD SUMF1-GT (Primary) mice (mean = 3.02, SD = 0.01) and MSD SUMF1-GT (Secondary) mice (mean = 3.27, SD = 0.11) as compared to untreated WT mice (mean = 1.27, SD = 0.57) (**Figure 3F**).

Lastly, we examined the stability of our lentiviral vector construct in transplanted mice by measuring the vector copy numbers in bone marrow cells at 4 months post-transplant of both primary and secondary transplanted mice. By conducting ddPCR targeting the Psi region of our lentiviral backbone and normalizing the data to a housekeeping gene in mice, *Pcpb2*, we determined that the vector copy numbers remained within a clinically relevant range of 0.4 to 1.9 in both primary and secondary transplanted mice (**Figure 3G**). Together, these data suggest that *SUMF1* HSCT-GT does not affect hematopoiesis and is durable in MSD mice.

### *Ex vivo SUMF1* gene therapy improves GAG levels in all tissues, but only partially rescues ARSA activities

Once we established that our *SUMF1* HSCT-GT approach was safe and long-lasting, we moved forward to investigate the efficacy of this approach in treating the biochemical deficits of MSD mice. Mirroring the biochemistry in human MSD cells, the tissues of the Ser153Pro MSD mouse model exhibit severe, multisystemic deficiencies in sulfatase activities.^38^ To elucidate the effects of *SUMF1* HSCT-GT on these sulfatase activities *in vivo*, we measured ARSA activities in whole-tissue homogenates of the brain, heart, lung, liver, and spleen of treated MSD mice at 6-months of age (4 months post-transplant). At baseline, we found that ARSA activity was significantly decreased in all tissues of 6-month-old, untreated MSD mice compared to WT (**Figure 4**). Previous studies suggest that this significant reduction in sulfatase activities is apparent as early as 6-weeks of age in the Ser153Pro MSD mice compared to WT.^38^ We found that *SUMF1* HSCT-GT was able to significantly increase ARSA activity in the spleen of treated mice (**Figure 4E**). ARSA activity increased about 2-fold after treatment as compared to untreated MSD mice. In contrast, ARSA activities of brain, heart, lung and liver remained unchanged after treatment. These data suggest that *SUMF1* HSCT-GT can rescue sulfatase activities in tissues with higher degrees of HSC-derived engraftment, like the spleen. After HSCT-GT, only a small fraction of each organ is repopulated by donor-derived cells. Therefore, because we are measuring enzyme activity in whole-tissue homogenates, we may only be able to detect improvements in sulfatase activities at the whole-organ level when a significant proportion of donor cells are engrafted into an organ.

**Figure 4.**
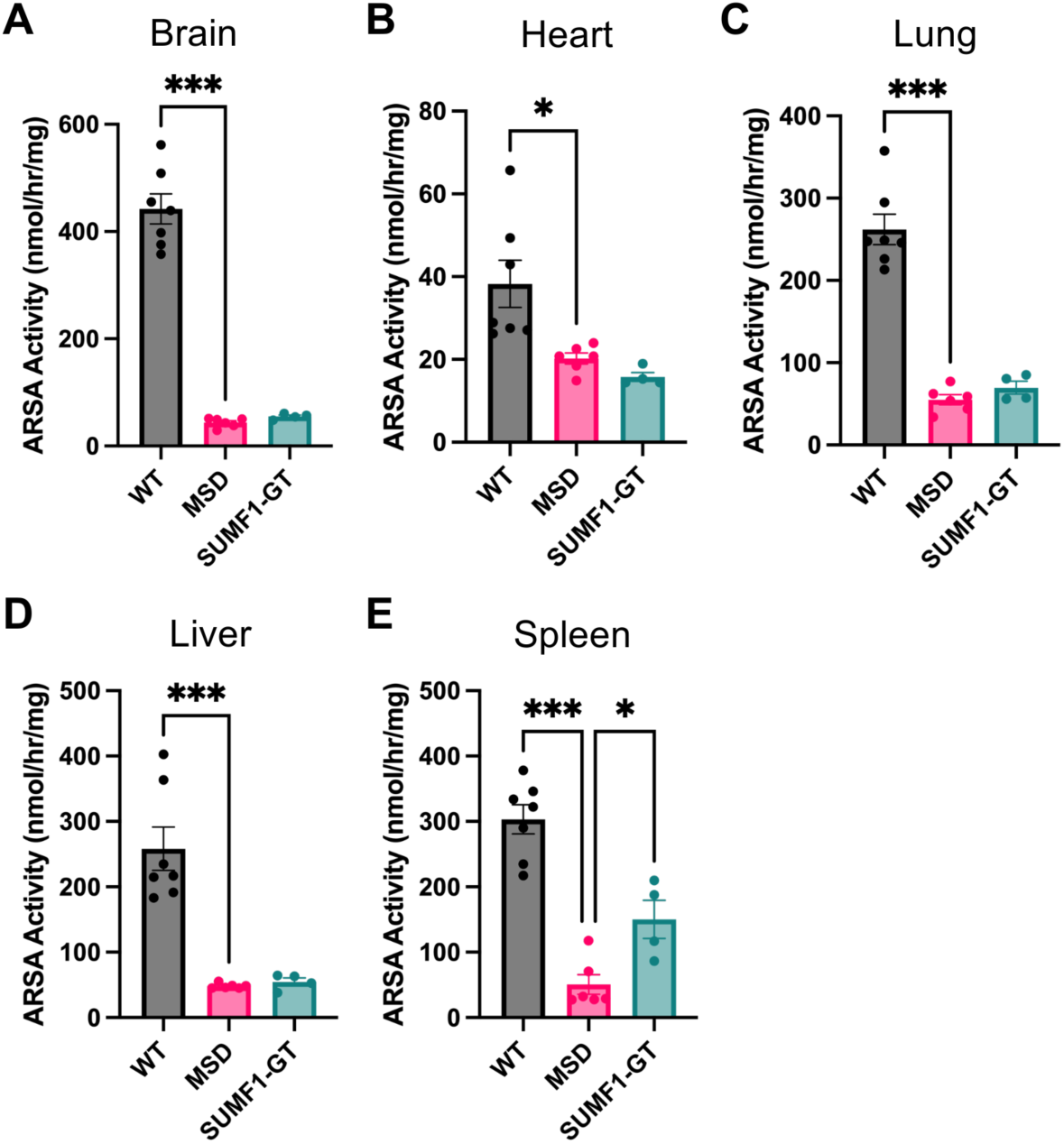
*SUMF1 ex vivo* gene therapy rescues ARSA activity in the spleen. ARSA activity quantifications (nmol/hr/mg total protein) for A) brain, B) heart, C) lung, D) liver, and E) spleen tissue homogenates from WT mice (WT), untreated MSD mice (MSD), or MSD mice receiving *SUMF1* HSCT-GT (SUMF1-GT). *N* = 4-7 mice, one-way ANOVA, followed by Bonferroni’s multiple comparisons test to compare each group to the untreated MSD control group. For all panels: *p < 0.05, ***p <0.001. Data are represented as mean ± SEM.

We next sought to evaluate the ability of *SUMF1* HSCT-GT to reduce the pathological storage of GAG subspecies in MSD mice. As with human MSD cells, the Ser153Pro MSD mouse model exhibits accumulation of overall GAGs in the liver and kidney, as measured by Alcian blue staining.^38^ To quantify the levels of GAG subspecies more specifically, we performed mass spectrometry using the endogenous, NRE biomarker method on whole-tissue homogenates of mouse brain, heart, lung, liver, and spleen at 6 months of age.^36,37^ For all tissue types, untreated Ser153Pro MSD mice showed accumulation of GAG subspecies associated with MPS II (UA-HN-UA-2S, UA-HNAc-UA-1S #1, and UA-HNAc-1S #2), MPS IIIA (HN-S, HN-UA-1S, HN-UA-HNAc-2S, HN-UA-HNAc-UA-1S, and HN-UA-HNAc-UA-2S), MPS IIID (GlcNAc-6s and HNAc(S)-UA #1) and MPS IVA/VI (GalNAc-6S, HNAc(S)-UA #2, HNAc-UA-HNAc-2S) at baseline as compared to WT mice (**Figure 5**).

**Figure 5.**
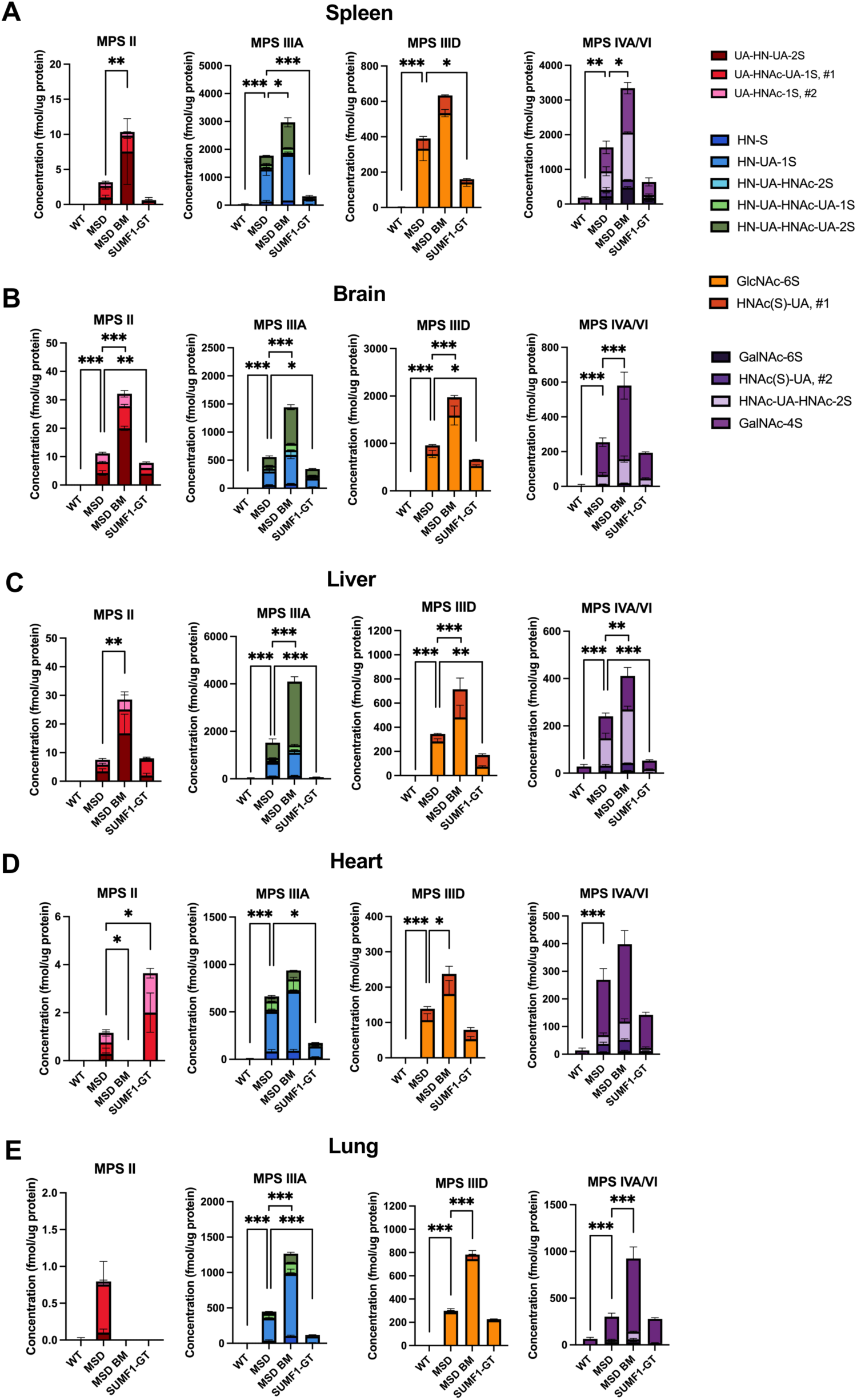
*SUMF1 ex vivo* gene therapy reduces GAG accumulation, while transplant alone exacerbates it *in vivo*. Quantification of GAG subspecies associated with MPS II, IIIA, IIID, and IVA/VI of untreated WT mice (WT), untreated MSD mice (MSD), MSD mice receiving non-transduced MSD bone marrow (MSD BM), and MSD mice receiving *SUMF1* HSCT-GT (SUMF1-GT) for A) spleen, B) brain, C) liver, D) heart, and E) lung tissue homogenates. Concentrations of GAGs (fmol/μg protein) were normalized to μg of total protein. n = 2-7 mice. Data are represented as mean ± SEM. For all panels: *p < 0.05, **p < 0.01, ***p < 0.001.

To determine the effects of the transplant process itself on MSD mice, we first performed HSCT experiments where we transplanted untreated MSD lineage-negative donor HSCs into naïve, irradiated MSD recipient mice as controls. We then measured the levels of GAG subspecies in the brain, heart, lung, liver and spleen of these mice at 6 months of age (4 months post-transplant). Interestingly, control MSD mice that received non-transduced MSD HSCs (MSD BM) showed a dramatic increase in GAG levels as compared to untreated MSD mice (**Figure 5**). Specifically, the accumulation of GAG subspecies associated with MPS IIIA, IIID, and IVA/VI worsened in MSD BM control mice in all tissues as compared to untreated MSD mice. The abundance of MPS II GAG subspecies in the heart and lung were at the lower level of detection and thus levels were unable to be compared between cohorts. We found that our *SUMF1* lentiviral vector (SUMF1-GT) was able to significantly rescue the GAG accumulation associated with the transplant process (**Figure 5**). The spleen, brain, liver, heart, and lung of *SUMF1* HSCT-GT-treated mice demonstrated improvement of storage for all GAG subspecies compared to both MSD BM control and untreated MSD mice.

### Brain microgliosis is reduced in mice treated with *ex vivo SUMF1* gene therapy

The Ser153Pro MSD mouse model has been shown to recapitulate human phenotypes and exhibit neuroinflammation in the brain including microgliosis and astrocytosis.^38^ *SUMF1* HSCT-GT has the potential to rescue MSD brain phenotypes as corrected, transplanted HSCs can repopulate bone-marrow derived cell populations, including microglia. In our therapeutic approach, HSCs transduced with our *SUMF1* lentiviral vector could cross the blood-brain barrier after transplant and differentiate into resident microglia-like cells to replace diseased microglia.

To study the effects of *SUMF1* HSCT-GT on brain microgliosis in the Ser153Pro MSD mouse model, we performed immunohistochemical analyses of ionized calcium-binding adaptor molecule 1 (Iba1)-labeled cells in the cortex of treated mice at 6-months of age (4-months post-transplant). Iba1 is a calcium-binding protein involved in membrane ruffling and phagocytosis of microglia, making it a classical marker of activated microglia. At baseline, 6-month-old untreated MSD mice showed significant increases in Iba1 staining compared to WT mice, as measured by fluorescence intensity and cell counting (**Figure 6**). These findings confirm that Ser153Pro MSD mice exhibit microgliosis in the cortex. We found that *SUMF1* HSCT-GT reduced this brain microgliosis in MSD mice, as indicated by significant decreases in Iba1 fluorescence intensity and cell counts in cortical sections of treated mice compared to untreated mice (**Figure 6**).

**Figure 6.**
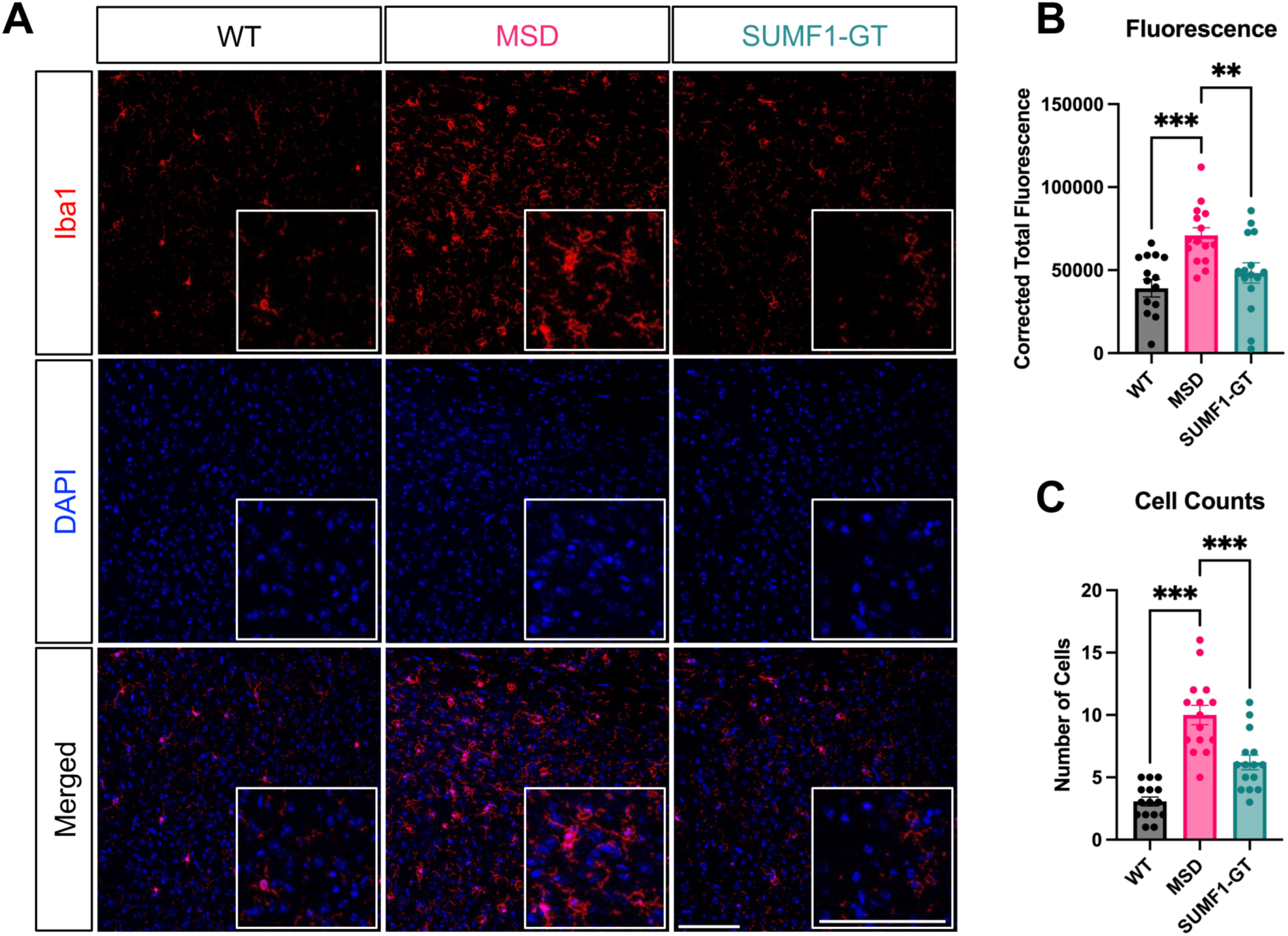
*SUMF1 ex vivo* gene therapy rescues microgliosis in the cortex. A) Representative immunofluorescence images of Iba1+ cells in cortical brain sections of WT mice (WT), untreated MSD mice (MSD), and MSD mice receiving *SUMF1* HSCT-GT (SUMF1-GT) reveal decreased microgliosis after treatment. Labeling with anti-Iba1 antibody (red fluorescence) and DAPI (nuclei, blue). Scale bars = 100 μm. B) Quantification of Iba1 immunofluorescence intensity confirms significant decrease in *SUMF1* HSCT-GT animals. C) Quantification of Iba1+ cells in cortical sections validates decreased microgliosis in treated mice. For all quantifications, three biological replicates were analyzed with 5 images per replicate, one-way ANOVA, followed by Bonferroni’s multiple comparisons test to compare each group to the untreated MSD control group. For all panels: **p < 0.01, ***p <0.001. Data are represented as mean ± SEM.

In addition, we examined the ability of *SUMF1* HSCT-GT to rescue brain astrocytosis in the Ser153Pro MSD mice by analyzing the abundance of glial fibrillary acidic protein (GFAP)-labeled cells in the cortex of treated mice at 4 months post-transplant. GFAP is an intermediate filament and a significant component of the astrocyte cytoskeleton, enabling us to assess levels of astrocytosis in the brain. We confirmed previous findings that untreated Ser153Pro MSD mice demonstrate high levels of astrocytosis in the cortex compared to WT mice by measuring GFAP fluorescence intensity and cell counts (**Figure S1**). However, MSD mice treated with *SUMF1* HSCT-GT did not show reduced levels of GFAP fluorescence intensity or cell counts in the cortex compared to untreated mice. Overall, these data suggest that our *SUMF1* HSCT-GT approach does not rescue astrocytosis but significantly improves microgliosis in MSD mice.

### *SUMF1* gene therapy only partially restores motor function, but normalizes neurocognitive and neurodegenerative behavioral phenotypes in MSD mice

Finally, we investigated the functional effects of *SUMF1* HSCT-GT in MSD mice by evaluating the rescue of behavioral phenotypes, as there is not an apparent survival phenotype in this MSD model.^38^ To evaluate potential preclinical endpoints, we conducted an initial battery of behavioral tests on untreated mice to characterize neuromuscular phenotypes in this new MSD mouse model.

Open field and elevated zero testing results did not differ between 6-month-old MSD and WT mice (**Figure S2**). Conversely, rotarod and grip strength testing revealed significant motor function impairments in untreated Ser153Pro MSD mice compared to WT at 6-months of age (**Figures 7A and 7B**). The rotarod assessment is a classic test of motor coordination and balance in which mice must continuously walk on a rotating rod that increases speed without falling off. WT mice perform significantly better on the rotarod each day than untreated MSD mice, staying on for 150-200 seconds and 100 seconds, respectively (**Figure 7A**). These data demonstrate that Ser153Pro MSD mice exhibit deficits in motor functioning. However, we found that both control MSD mice that received non-transduced MSD bone marrow (MSD BM) and our experimental MSD mice that received *SUMF1* HSCT-GT (SUMF1-GT) did not exhibit significant improvements in the rotarod test (**Figure 7A**). Control MSD BM mice stayed on the rotarod for about 150 seconds per day while SUMF1-GT mice only stayed on for about 125 seconds per day, neither statistically significant from untreated MSD mice (**Figure 7A**).

**Figure 7.**
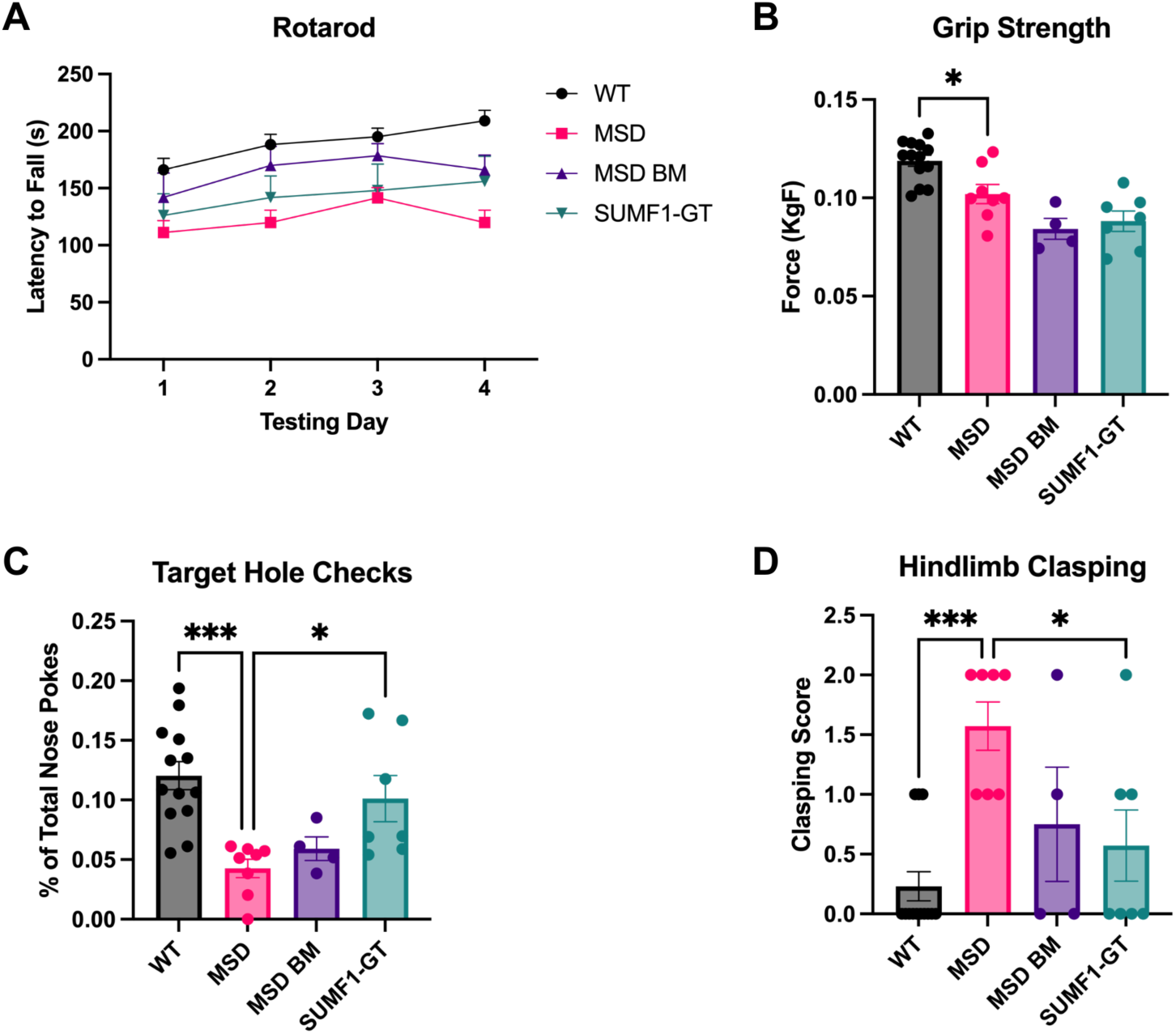
*SUMF1 ex vivo* gene therapy improves cognitive function and neurodegeneration, but not motor phenotypes. A) Quantification of rotarod assay for WT mice (WT), untreated MSD mice (MSD), MSD mice receiving non-transduced MSD bone marrow (MSD BM), and MSD mice receiving *SUMF1* HSCT-GT (SUMF1-GT) suggests partial improvement in motor coordination phenotypes after treatment. The latency (seconds) of the mouse to fall off the rotarod was measured over 4 days. *N* = 4-11 mice, one-way ANOVA at each testing day, followed by Bonferroni’s multiple comparisons test to compare each group to the untreated MSD control group. Control MSD BM and experimental SUMF1-GT groups were not statistically significant from the untreated MSD group at any time point. Data are represented as mean ± SEM. B) Quantification of grip strength assay for WT mice (WT), untreated MSD mice (MSD), MSD mice receiving non-transduced MSD bone marrow (MSD BM), and MSD mice receiving *SUMF1* HSCT-GT (SUMF1-GT) reveals no muscular improvement after treatment. The force (KgF) generated by the mouse forelimb is reported. *N* = 4-13 mice, one-way ANOVA, followed by Bonferroni’s multiple comparisons test to compare each group to the untreated MSD control group. Data are represented as mean ± SEM. C) Quantification of Barnes maze assay for WT mice (WT), untreated MSD mice (MSD), MSD mice receiving non-transduced MSD bone marrow (MSD BM), and MSD mice receiving *SUMF1* HSCT-GT (SUMF1-GT) shows improved neurocognitive function after *SUMF1* HSCT-GT. Percentage (%) of total nose pokes is the number of times an animal checked the target hole divided by the total number of times it checked any hole. *N* = 4-13 mice, one-way ANOVA, followed by Bonferroni’s multiple comparisons test to compare each group to the untreated MSD control group. Data are represented as mean ± SEM. D) Quantification of hindlimb clasping scores for WT mice (WT), untreated MSD mice (MSD), MSD mice receiving non-transduced MSD bone marrow (MSD BM), and MSD mice receiving *SUMF1* HSCT-GT (SUMF1-GT) reveals rescue of neurodegenerative phenotype after *SUMF1* HSCT-GT. *N* = 4-11 mice, one-way ANOVA, followed by Bonferroni’s multiple comparisons test to compare each group to the untreated MSD control group. Data are represented as mean ± SEM. For all panels: *p < 0.05, **p < 0.01, ***p <0.001.

Additionally, we performed forelimb grip strength assessments on untreated and treated Ser153Pro MSD mice to characterize differences in muscular strength. The grip strength test measures the maximum force generated by the forelimb muscles. WT mice are significantly stronger than MSD mice, as indicated by the increase in forelimb force (KgF) generated by WT mice (**Figure 7B**). However, we again found that neither MSD BM controls nor SUMF1-GT mice exhibited increased forelimb force, suggesting that these approaches are not able to rescue muscular deficiencies in MSD mice. The data suggest that our *SUMF1* HSCT-GT approach only partially rescues motor coordination and strength.

We next evaluated cognitive function by conducting Barnes maze assays on treated and untreated Ser153Pro MSD mice. The Barnes maze test is an assay that measures spatial learning and memory, in which mice are trained to learn a target hole and later tested to identify the correct hole. On the testing day, the proportion of times that WT mice identify the target hole is significantly greater than that of untreated MSD mice (**Figure 7C**). This suggests that WT mice correctly learn and remember the target hole while MSD mice do not, reflecting the cognitive deficits of MSD mice. Control MSD mice receiving non-transduced bone marrow (MSD BM) do not show significant improvements on the Barnes maze, indicating that the transplant process itself is insufficient to rescue this cognitive phenotype. Conversely, MSD mice treated with *SUMF1* HSCT-GT (SUMF1-GT) performed significantly better than untreated MSD mice, correctly identifying the target hole more often. These findings suggest that *SUMF1* HSCT-GT effectively rescues the spatial learning and memory deficits in MSD mice.

Finally, we examined neurodegeneration in the Ser153Pro MSD mouse model by assessing for a hindlimb clasping phenotype. Mice with higher scores on the hindlimb clasping test demonstrate more severe neurodegeneration and disease progression, as established previously.^40^ These assays confirmed that Ser153Pro MSD mice have a severe neurodegeneration phenotype, as indicated by the significantly higher clasping score than WT mice (**Figure 7D**). Paralleling the neurocognitive rescue seen with the Barnes maze, only mice treated with *SUMF1* HSCT-GT exhibited rescue of this neurodegeneration phenotype, as demonstrated by a significantly decreased clasping score compared to untreated MSD mice. Overall, these behavioral tests suggest that while *SUMF1* HSCT-GT only partially restores motor function, it successfully improves neurocognitive function and measures of neurodegenerative disease progression in MSD mice.

## DISCUSSION

MSD is an untreatable, progressive, lethal disorder. Patients suffer from significant morbidity, including multi-system organ dysfunction and neurologic regression. Here, we investigated if *SUMF1 ex vivo* gene therapy with hematopoietic stem cell transplantation could improve disease phenotypes in MSD models. We found that *SUMF1* HSCT-GT improved storage accumulation in several affected organs of an MSD mouse model, including the spleen, liver, lungs, heart, and brain. In addition, this therapeutic approach reduced neuroinflammation and improved neurocognitive function.

In designing this work, we prioritized a study design that would allow us to evaluate the effect of *SUMF1* HSCT-GT in a more clinically relevant manner. Specifically, we 1) chose an MSD mouse model that shares genetic features with the human disease, 2) treated mice at an age that may be clinically translatable to human patients, and 3) selected preclinical study endpoints that are similar to the biochemical phenotypes and neuromuscular deficits seen in MSD patients.

The recently-created *Sumf1* p.Ser153Pro mouse model^38^ utilized here models a severe, recurrent human pathogenic variant (human variant *SUMF1* p.Ser155Pro). Patients with MSD who survive beyond the neonatal period typically carry at least one missense variant, such as the Ser155Pro allele, as biallelic stop-gain variants are universally lethal shortly after birth.

Similarly, the *Sumf1-/-* knockout mouse dies in the neonatal period. The Ser153Pro human mutation mouse model better recapitulates human disease and allowed us to evaluate the efficacy of a post-natal MSD therapeutic.

Because of its rarity and the lack of MSD newborn screening, most patients with MSD are diagnosed months to years after symptom onset.^14^ Therefore, new therapies must improve outcomes in symptomatic animal models and patients. We modelled this phenomenon by treating adult (8-week-old) MSD mice with *SUMF1* HSCT-GT. This is in contrast to many studies of gene therapy in neuronopathic LSDs, where dosing occurs in the neonatal period before significant storage accumulation has occurred. At 8-weeks of age, Ser153Pro MSD mice already show poor growth and GAG accumulation in multiple organs.^38^ For example, significant GAG accumulation is detected in the affected mice as early as 6-weeks of age.^38^ Encouragingly, *SUMF1* HSCT-GT improved many disease phenotypes in the mice, despite dosing at a later time point. Furthermore, the rescue of GAG accumulation suggests that post-symptomatic treatment with our *SUMF1* HSCT-GT approach effectively reverses storage phenotypes in MSD mice. Earlier dosing strategies will likely provide even more benefit in MSD. Early identification of MSD patients may soon be feasible, as MSD cases will likely be identified through emerging newborn screening programs for single sulfatase disorders^41^ such as MLD and MPS II.

While it is encouraging that our *SUMF1* HSCT-GT improves storage accumulation and neuroinflammation, it is important that this approach also augments behavioral phenotypes in the mice. When selecting functional outcome measures in this preclinical study, we focused on endpoints that represent neurological manifestations of disease, as these are a major cause of morbidity in patients. We identified novel behavioral phenotypes in the Ser153Pro mouse model that capture both neurocognitive dysfunction and CNS neurodegeneration, key features of the human disorder. Ser153Pro mice show poor cognitive function as measured by the Barnes maze assay,^42^ and potential atrophy and disruption to the cerebello-cortico-reticular pathway as measured by a hindlimb clasping phenotype.^40,43^ Importantly, *SUMF1* HSCT-GT corrected both of these functional phenotypes in MSD mice.

As any translational program moves towards clinical trials for MSD, it will be essential to evaluate the durability of biochemical rescue as measured by tractable, disease-relevant therapeutic biomarkers. GAG accumulation has been proposed as an MSD biomarker; however, a previous study of the Ser153Pro mouse did not demonstrate GAG elevations in the brain through nonspecific, Alcian blue staining.^38^ We recently completed a comprehensive human patient biomarker study using a more-specific, mass spectrometry-based GAG detection method.^37,44^ We found that endogenous GAG nonreducing end subspecies that accumulate in MPS IIIA, MPS IIID and MPS II better correlate with disease severity in urine samples from MSD patients, than other subspecies.^45^ Here, we found that the Ser153Pro mouse accumulates these same specific GAG subspecies in the brain and most other organs examined, and *SUMF1* HSCT-GT improved these clinically relevant biomarkers in the spleen, brain, liver, heart, and lung of treated MSD mice.

Although we observed a benefit of *SUMF1* HSCT-GT in our MSD mouse model, further studies are needed to build on this proof-of-concept study and optimize this approach. One limitation that could be improved upon is the plateau in efficacy that we see in our *in vitro* system at higher vector copy numbers. We found that increasing protein levels of the sulfatase activating factor, FGE, through gene therapy only increased downstream target sulfatase activation to a certain degree, until a plateau was reached (**Figures 1D-F**). This is likely due to a limited amount of available downstream target sulfatases. Future studies could investigate ways of increasing the levels of available downstream sulfatases, in addition to replacing the missing FGE activity through gene therapy. Ultimately, this may be required to optimize clinical efficacy in MSD models and patients. Optimization of FGE and downstream sulfatase dosing may facilitate a more robust rescue of motor phenotypes, as our current therapy did not significantly improve these measures of disease.

A transformative therapy for MSD must be safe, in addition to being efficacious. All transplanted mice lived until the study endpoint (4 months after transplantation, 6 months of age) without any signs of transplant-related morbidity or deficits in normal hematopoiesis. All mice treated with *SUMF1* HSCT-GT achieved high engraftment levels (88-100%) that were durable for up to 8 months post-transplant (i.e., 4 months post primary transplant, followed by 4 months of secondary transplant). Flow cytometric analysis demonstrated no gross abnormalities of erythroid or lymphoid populations in HSCT-GT treated animals.

As HSCT-GT uses an autologous stem cell source, as opposed to standard allogenic transplant, there is a reduced risk of immune complications such as graft-vs-host disease and graft rejection. However, HSCT-GT presents a potential genotoxicity risk given that lentiviral vectors integrate into the genome.^46^ Our *SUMF1*-expressing lentiviral vector has been engineered to minimize the risk of insertional oncogenesis. Specifically, insulator elements were included in the vector to minimize undesirable promoter activation and subsequent aberrant transcription at sites of vector integration.^32,47–49^ The vector backbone used here has been previously evaluated through *in vitro* oncogenesis assays and has not demonstrated induced clonal expansion of transduced mouse HSCs (see the companion paper submitted by Tricoli et al). Expanded safety and toxicity studies of HSCT-GT in MSD mice and large animal models will be needed to ensure the safety of this therapeutic approach further.

Another key strategy to minimize potential cancer risk after HSCT-GT is to minimize the vector copy number (VCN) required to achieve a therapeutic benefit. A recent meta-analysis of data from all available *ex vivo* gene therapy trials for monogenic disorders found that the mean VCN achieved across all trials was 1.6 (range 0.05-9.4).^50^ The improvements observed in our MSD mice after HSCT-GT were achieved with a VCN of 0.4-1.9, which is well within the range for what is considered safe in patients.

Multiple alternative therapeutic strategies are being developed for MSD. Recently, two MSD patients underwent HSCT, a first for MSD. Full outcomes are forthcoming, but both patients tolerated the procedure well.^51^ A study of HSCT in an MSD mouse model was conducted in parallel. Preliminary reports show robust engraftment of donor cells, and no procedure-induced toxicity.^52^ While these efforts are promising, unfortunately, HSCT alone is often insufficient to stop the neurologic regression that occurs in several related lysosomal storage disorders.^23^

CNS-directed adeno-associated virus (AAV)-mediated *SUMF1* gene replacement is also in preclinical development for MSD.^53,54^ While brain-directed AAV gene therapy has shown promise in several preclinical models of neuronopathic lysosomal storage disorders, it may not address many somatic manifestations of MSD, such as orthopedic complications.^12,14^ Currently-utilized AAV capsids have a limited ability to target bone and cartilage. However, unlike AAV-based therapies, recent data in Hurler syndrome (MPS I) and Hunter syndrome (MPS II) suggest that HSCT-GT may ameliorate bone phenotypes.^28,30^

Finally, small molecule strategies are also in development for MSD. A recent drug repurposing screen identified two retinoids, tazarotene and bexarotene, as potential therapies in MSD.^35^ Both drugs can increase sulfatase activities and reduce disease phenotypes in cells from MSD patients. Future studies are needed to compare the efficacy of these therapeutic modalities alone, and in combination, in both preclinical models and MSD patients.

In conclusion, *ex vivo* lentiviral-mediated *SUMF1* gene therapy with HSCT improves biochemical and functional markers of disease in a clinically relevant MSD mouse model. Additional safety and efficacy studies are needed to advance this approach closer to the clinic. Ultimately, this foundational work could support first-in-human clinical trials to improve somatic symptoms and neurologic outcomes for patients living with MSD.

## MATERIALS AND METHODS

### Cell lines

Previously described^35^ immortalized WT fibroblasts and immortalized fibroblasts from an MSD patient homozygous for the severe, pathogenic *SUMF1* c.463C>T (p.Ser155Pro) variant (MSD) were grown in cell culture at 37°C under 5% CO_2_ in Dulbecco’s Modified Eagle’s Medium (Corning, Inc., Corning, NY) containing 10% HyClone Characterized Fetal Bovine Serum (Cytiva, Marlborough, MA) and 1% penicillin-streptomycin (ThermoFisher Scientific, Waltham, MA). The cells were regularly tested to exclude mycoplasma contamination.

### Lentiviral plasmid and virus generation

All lentiviral constructs were designed in-house, and all plasmids were manufactured by Genscript (Piscataway, NJ). Plasmid DNA was expanded and isolated using a QIAGEN Plasmid Maxi Kit (Germantown, MD). The lentiviral vectors were assembled using a 3rd generation lentiviral system and CalFectin^TM^ Mammalian DNA Transfection Reagent (SignaGen Laboratories, Rockville, MD) in 293T cells according to the manufacturer protocol. Briefly, 5 mL of complete Iscove’s Modification of Dulbecco’s Modified Eagle Medium (Corning, Corning, NY) with 10% HyClone Characterized Fetal Bovine Serum (Cytiva, Malborough, MA) and 1% penicillin-streptomycin (ThermoFisher Scientific, Waltham, MA) was added to each 10 cm dish of 90% confluent 293T cells 60 minutes before transfection. For each dish, 3.7 μg of pMDLg/RRE DNA, 2.2 μg of pMD2.G DNA, and 1.83 μg of pRSV-Rev DNA (Addgene, Watertown, MA) in addition to 7.32 μg of lentiviral transfer vector DNA in 500 uL of serum-free Dulbecco’s Modified Eagle Medium with High Glucose (Corning, Corning, NY) were added to CalFectin Reagent diluted in an equal volume of serum-free Dulbecco’s Modified Eagle Medium. The CalFectin-DNA complexes were incubated for 15 minutes at room temperature before being added dropwise onto the medium of each 293T dish. The next day, fresh complete Iscove’s Modification of Dulbecco’s Modified Eagle Medium (Corning, Corning, NY) with 10% HyClone Characterized Fetal Bovine Serum (Cytiva, Malborough, MA) and 1% penicillin-streptomycin (ThermoFisher Scientific, Waltham, MA) was added to each 10 cm dish of 293T cells. The viral supernatant was collected 48 hours post-transfection and concentrated via ultracentrifugation.

### Mouse lines, HSC lineage negative isolation, and bone marrow transplantation

The Sumf1(S153P) mice (stock number 31558, C57BL/6J-*Sumf1*^em8Lutzy^/Mmjax, The Jackson Laboratory, Bar Harbor, ME) were maintained as homozygous colony at The Children’s Hospital of Philadelphia animal facility. Lineage-negative HSCs were isolated from the bone marrow of donor mice via immunomagnetic separation using a mouse lineage cell depletion kit (Miltenyi Biotec, Auburn, CA). Lineage-negative HSCs were transduced overnight at a viral multiplicity of infection (MOI) of 2-50 in Dulbecco’s Modified Eagle Medium (Corning, Corning, NY) supplemented with 10% HyClone Characterized Fetal Bovine Serum (Cytiva, Malborough, MA), 1% penicillin-streptomycin (ThermoFisher Scientific, Waltham, MA), 100 ng/mL recombinant murine stem cell factor (SCF), 10 ng/mL recombinant murine interleukin-6 (IL-6), 6 ng/mL interleukin-3 (IL-3) (PeproTech, Rocky Hill, NJ) and 2 uL Lentiblast Premium Transduction Enhancer (OZBiosciences, San Diego, CA). Before receiving lineage-negative HSCs, recipient mice (8-weeks old) were irradiated by receiving two rounds of 500 cGy using an X-ray irradiator 4 hours apart. 500,000-1,000,000 viable lineage-negative donor HSCs in phosphate-buffered saline (PBS) (Gibco, Thermo Fisher Scientific, Waltham, MA) were then injected retro-orbitally into irradiated recipient mice of the opposite sex.

### Western blotting

Total protein concentrations were measured in cell lysates using the Pierce BCA Protein Assay Kit (ThermoFisher Scientific, Waltham, MA). A total of 30 μg of protein fraction for each sample was separated by electrophoresis using NuPAGE^TM^ 4-12% Bis-Tris Protein Gel (ThermoFisher Scientific, Waltham, MA). Proteins were transferred to a polyvinylidene difluoride (PVDF) membrane using an XCell SureLock Mini-Cell Electrophoresis System (ThermoFisher Scientific, Waltham, MA). The membrane was blocked using 5% milk in Tris-Buffered Saline (Bio Basic, Markham, Ontario, Canada) with Tween-20 for 45 minutes at room temperature. After briefly rinsing with Tris-Buffered Saline (Bio Basic, Markham, Ontario, Canada) with Tween-20, the membrane was incubated overnight at 4°C with one of the following antibodies: FGE (R&D Systems, MAB2779, Minneapolis, MN; dilution 1:1000) or Vinculin (Abcam, ab91459, Cambridge, UK; dilution 1:2000). The membrane was then washed with Tris-Buffered Saline with Tween-20 and incubated in one of the following secondary antibodies: Goat Anti-Rabbit IgG H&L (HRP) (Abcam, ab6721, Cambridge, UK; dilution 1:10,000) or Goat Anti-Mouse IgG H&L (HRP) (Abcam, ab205719, Cambridge, UK; dilution 1:10,000) for 1.5 hours at room temperature. Clarity Max^TM^ Western ECL Substrate (BioRad, Hercules, CA) and ChemiDoc XRS+ Imaging System (BioRad, Hercules, CA) were used for visualization.

### Sulfatase activity assays

ARSA and ARSB activities were quantified using previously published protocols.^55^ Briefly, 50 μg of protein extract of whole cell lysates was incubated at 37°C for 1 hr with the artificial p-nitrocatecholsulfate substrate (Santa Cruz Biotechnology, Dallas, TX, sc-238927A) and either sodium pyrophosphate (for ARSA) or barium acetate (for ARSB). For ARSA activity of tissues, 100 μg of tissue homogenate was incubated at 37°C for 1 hr with the artificial p-nitrocatecholsulfate substrate (Santa Cruz Biotechnology, Dallas, TX, sc-238927A) and sodium pyrophosphate. The absorbance was measured at 515 nm in a BioTek Epoch Microplate Spectrophotometer (Agilent, Santa Clara, CA). ARSA and ARSB units are defined as the amount of 4-nitrocatechol liberated in 1 hr per mg of total protein. SGSH activity was measured from whole cell lysates using a previously published two-step protocol^56^ and using 4MU-αGlcNS (Chem-Impex Int’l Inc, Wood Dale, IL, 30694) as fluorogenic substrate and with 4-methylumbelliferone (Sigma Aldrich, St. Louis, MO) as standard. A unit of SGSH activity is defined as the amount that will liberate 1 nmol 4-MU at 37°C in 17 hr.

### Glycosaminoglycan quantification

GAG fragments (i.e., sulfated oligosaccharides) in cells and tissues were measured using a modified protocol as described.^36,37^ Chondroitin disaccharide di-4S (dUA-GalNAc-4S, Biosynth, Staad, Switzerland) was used as the internal standard (IS). Cell lysates and tissue homogenates equivalent to 100 µg protein were used. The UPLC-MS/MS analysis was conducted on a Waters TQ-S triple quadrupole mass spectrometry coupled to an ACQUITY Classic UPLC system. Three previously reported markers, HexN-UA-1S (MPS IIIA), GlcNAc-6S (MPS IIID), and GalNAc-4S (MPS VI), were analyzed.^36,37^ Two novel markers, UA-HexN-UA-2S (509.8 -> 422.7) and HexN-UA-HexNAc-UA-2S (611.4 -> 524.1) associated with MPS I/II and MPS IIIA, respectively, were also included. The apparent concentration was calculated by multiplying the response ratio of the analyte and the IS by the amount of IS added (fmol), then dividing by the amount of protein (µg).

### Flow cytometry

Flow cytometry analysis was conducted with a BD FACSCalibur instrument (BD Biosciences, Franklin Lakes, NJ) to examine erythroid and lymphoid populations in the bone marrow. Specifically, FITC Anti-Mouse CD71 (113806) and APC Anti-Mouse Ter119 (116212), along with PE Anti-Human/Mouse CD44 (103008) antibodies (Biolegend, San Diego, CA), were utilized for the erythroid populations. PE/Cyanine7 anti-mouse/human CD45R/B220 Antibody (103222) and APC anti-mouse CD3 Antibody (100236) antibodies (Biolegend, San Diego, CA) were utilized for the lymphoid populations. Subsequently, the acquired data was processed and interpreted using FlowJo software developed by Tree Star Inc. (Ashland, OR).

### ddPCR

Primers/probes targeting the Psi region of the lentiviral backbone (BioRad Custom Assay, Hercules, CA) were used to detect viral integration: Primer-F 5’-ACCTGAAAGCGAAAGGGAAAC-3’; Primer-R 5’-CGCACCCATCTCTCTCCTTCT-3’; and FAM Probe 5’-AGCTCTCTCGACGCAGGACTCGGC-3’. Primers/probes targeting the male-specific mouse *Zfy1* gene (BioRad Custom Assay, Hercules, CA) were used to detect engraftment as opposite sex donors/recipients were used: Primer-F 5’-AAGTGCTTCTTGGCATCACC-3’; Primer-R 5’-ATCCAAAGACTGCTCCACCT-3’; and FAM Probe 5’-TGGGTTTGGTGTCTGCAGATGGT-3’. Primers/probes to detect mouse *Pcbp2* (BioRad Custom Assay, Hercules, CA) were used as an endogenous reference in mouse samples: Primer-F 5’-CCAGTCTGCTTGGCATGAAA-3’; Primer-R ‘5-GGTACCCTTAGCAGCAGACA-3’; and HEX Probe 5’-CCCATCCCTCTCCTGGCTCTAA-3’. Primers/probes to detect human *RPP30* (BioRad Custom Assay, Hercules, CA) were used as an endogenous reference in human samples: Primer-F 5’-ACCACCACATCCCAGCTAAT-3’; Primer-R 5’-GACTGCTTGAATCTGCCAGG-3’; and HEX Probe 5’-TGCCATGTCCAGGCTAGTCTCAAA-3’. ddPCR reactions were conducted with ddPCR Supermix for probes (no dUTP) (BioRad, Hercules, CA). ddPCR plates were prepared using a BioRad automated droplet generator and Px1 plate sealer (Hercules, CA). The PCR was conducted on a Bio-Rad C1000 thermocycler and analyzed on the Bio-Rad QX200 (Hercules, CA).

### Immunofluorescence

Brains were harvested from 6-month-old mice under isoflurane anesthesia. Brains were fixed via transcardial perfusion with PBS followed by 4% paraformaldehyde. Brain tissue was then post-fixed in 4% paraformaldehyde for 24 hours and cryopreserved in 30% sucrose for 72 hours at 4°C. Brains were embedded in optimal cutting temperature compound (OCT; Sakura Finetek, Japan, 4583) and stored at -80°C until use. Tissue was cryo-sectioned into 20 μm serial sections and affixed to slides for immunohistochemistry.

For immunofluorescence staining, slides were permeabilized using 0.1% Triton X-100 (Sigma-Aldrich, Burlington, MA) diluted in PBS for 10 minutes at room temperature. Slides were then blocked at room temperature using 10% Normal Goat Serum (Millipore, Burlington, MA) diluted in 0.1% Triton X-100 for 1.5 hours. Primary antibodies were diluted in blocking solution (10% Normal Goat Serum in 0.1% Triton X-100) and added to slides for incubation overnight at 4°C in a dark box. The following primary antibodies were used: Mouse Anti-NeuN Antibody (Millipore, Burlington MA, MAB377, dilution 1:100), Polyclonal Rabbit Anti-Glial Fibrillary Acidic Protein (Dako, Denmark, Z0334, dilution 1:200), Rabbit Anti-Iba1(Fujifilm, Japan, 019-19741, dilution 1:500). After removing primary antibodies and washing slides with PBS, secondary antibodies were diluted with NucBlue Fixed Cell Stain ReadyProbes (Invitrogen, Waltham MA, R37606, 1 drop per 1 mL) in blocking solution and incubated on slides for 30 minutes at room temperature in a dark box. The following secondary antibodies were used: Anti-Mouse IgG (H+L) Fab2 Alexa Fluor 488 Conjugate (Cell Signaling Technologies, Danvers MA, 4408S, dilution 1:50), Anti-Rabbit IgG (H+L) Fab2 Alexa Fluor 594 Conjugate (Cell Signaling Technologies, Danvers MA, 8889S, dilution 1:50 for GFAP, dilution 1:1000 for Iba1). Slides were washed three times with PBS after secondary antibody incubation. TrueBlack Lipofuscin Autofluorescence Quencher (Biotium, Fremont, CA) diluted in 70% ethanol was added to each slice for 30 seconds. The slides were then washed three times with PBS. Coverslips were mounted on slides using Prolong Gold mounting media (Thermo Scientific, Waltham, MA, P36930). Slides were cured for 2-3 hours at room temperature while shielded from light, then moved to 4°C for storage in the dark. Images for quantification were acquired using a Keyence BZ-X800 (Itasca, IL). Representative images were acquired using a Leica STELLARIS 5 Confocal Microscope (Wetzlar, Germany).

### Behavioral tests

For rotarod testing, mice were acclimated to the testing room for 30 minutes before start. An Ugo-Basile accelerating rotarod (Gemonio VA, Italy, model 47600) for mice was used for testing. On Day 1 of testing, mice were first subjected to a training period in which they walked on the rotarod at 5 rpm for 5 min. On Days 1-4, the mice were subjected to three trials, with 30-minute inter-trial rest periods. Each trial consists of the rotarod starting at 5 rpm and accelerating to 40 rpm over the course of 240 seconds. The time in seconds when the mouse fell from the rod was recorded and averaged for the three trials per mouse each day.

For grip strength testing, mice were acclimated to the testing room for 30 min before start. A Chatillon DFE-002 Force Gauge grip strength meter (Columbus Instruments, Columbus, OH) was used. The total peak force in kilogram-force (KgF) was measured five times per mouse with 30-minute inter-trial rest periods. For data analysis, the highest and lowest measurements were excluded for a total of three measurements per mouse. The mean of the three measurements per mouse was then quantified.

For Barnes maze testing,^42^ mice were acclimated to the testing room for 30 min before start. A Barnes maze table (San Diego Instruments, San Diego, CA. Cat #7001-0235) was used. On Days 1-4, mice were subjected to two training sessions with 20-minute inter-trial rest periods. During each training session, mice were allowed to explore the Barnes maze for 150 seconds. If the mouse entered the target hole before 150 seconds, the hole was covered to allow the mouse to habituate in the hole for 60 seconds. If the mouse did not enter the target hole before 150 seconds, the mouse was encouraged to enter the hole, and the hole was covered for 60 seconds. On Day 5, mice were subjected to one testing session in which the target hole was occluded, and the mouse was given 120 seconds to explore the Barnes Maze. The number of times the mouse checked the target hole on the test day divided by the total number of times the mouse checked any hole was quantified.

For the hindlimb clasping assay,^40^ mice were acclimated to the testing room for 30 minutes before start. Mice were lifted by the tail, with the ventral side towards the experimenter, and held for 5 seconds. If both mouse hindlimbs were fully extended outwards, the mouse received a score of 0. If one hindlimb was straight down with the paw clasped, the mouse received a score of 1. If two hindlimbs were straight down and the paws were clasped, the mouse received a score of 2. If the mouse formed a ball, the mouse received a score of 3.

Both male and female mice were used for these studies. For behavior and histology studies, the investigator was blinded to genotype when possible.

### Statistical analyses

All statistical analysis was performed using Prism (GraphPad software, San Diego, USA). Comparison of two independent groups was conducted by unpaired t-test. Comparison of three or more groups was conducted by one-way ANOVA, followed by Bonferroni’s multiple comparisons test to compare each group to the untreated WT control group (for flow cytometry analyses) or the untreated MSD control group (for all other analyses). Data were represented as mean ± standard error. Significance levels were displayed as follows: *p < 0.05, **p > 0.01, ***p < 0.001 for differences indicated in the figure legends. Differences were not statistically significant if not indicated.

### Study approval

All animal protocols were approved by the Institutional Animal Care and Use Committee at the Children’s Hospital of Philadelphia. Reporting of animal studies have been provided in accordance with ARRIVE guidelines.

## ACKNOWLEDGEMENTS

This work was funded by the Irish Health Research Board and MSD Action Foundation to R.C.A-N. and S.B.R. It is also supported by the CHOP Research Institute including The CHOP Gene Therapy for Inherited Metabolic Disorders Frontier Program (to R.C.A-N.), the Sickle Cell and Red Cell Disorders Curative Therapy Center (to S.B.R.) and the Acceleration-Seed program (to S.B.R.). Additional relevant funding includes NIH K23NS114113, U54TR002823 to L.A.A. We would like to thank Dr. Cathleen Lutz and Dr. Maximilliano Presa from the Jackson Laboratory for providing the mouse model. We also thank the MSD advocacy organizations and patients for their ongoing support and collaboration.

## AUTHOR CONTRIBUTIONS

Design of study: V.P., L.T., L.S., S.B.R., R.C.A-N.

Data acquisition and analysis of data: V.P., L.T., X.H., P.W., C.C.C., A.G., M.M.C., O.K., E.T., A.K., S.B.R., R.C.A-N.

Drafting of manuscript: V.P., L.T., X.H., C.C.C., M.M.C., R.C.A-N.

Critical review of data and manuscript: V.P., L.T., X.H., P.W., C.C.C., A.G., L.S., L.A.A., A.K., M.M.C., O.K., E.T., M.P., C.L., S.B.R., R.C.A-N.

## DECLARATION OF INTERESTS

RAN is an advisor to Latus Bio, AskBio, and Orchard Therapeutics. LA is an advisor to Takeda and Orchard Therapeutics. SR is a scientific advisory board member of Ionis Pharmaceuticals, Meira GTx, Vifor and Disc Medicine. He has been or is a consultant for GSK, BMS, Incyte, Cambridge Healthcare Res, Celgene Corporation, Catenion, First Manhattan Co., FORMA Therapeutics, Ghost Tree Capital, Keros Therapeutics, Noble insight, Protagonist Therapeutics, Sanofi Aventis U.S., Slingshot Insight, Spexis AG, Techspert.io, BVF Partners L.P., Rallybio, LLC, venBio Select LLC, ExpertConnect LLC, LifeSci Capital. No other authors report conflicts of interest.

## REFERENCES

1. Cappuccio, G., Alagia, M., and Brunetti-Pierri, N. (2020). A systematic cross-sectional survey of multiple sulfatase deficiency. Mol Genet Metab 130, 283–288. 10.1016/j.ymgme.2020.06.005.

2. Dierks, T., Schmidt, B., Borissenko, L.V., Peng, J., Preusser, A., Mariappan, M., and von Figura, K. (2003). Multiple sulfatase deficiency is caused by mutations in the gene encoding the human C(alpha)-formylglycine generating enzyme. Cell 113, 435–444. 10.1016/s0092-8674(03)00347-7.

3. Cosma, M.P., Pepe, S., Annunziata, I., Newbold, R.F., Grompe, M., Parenti, G., and Ballabio, A. (2003). The multiple sulfatase deficiency gene encodes an essential and limiting factor for the activity of sulfatases. Cell 113, 445–456. 10.1016/s0092-8674(03)00348-9.

4. Cosma, M.P., Pepe, S., Parenti, G., Settembre, C., Annunziata, I., Wade-Martins, R., Di Domenico, C., Di Natale, P., Mankad, A., Cox, B., et al. (2004). Molecular and functional analysis of SUMF1 mutations in multiple sulfatase deficiency. Hum Mutat 23, 576–581. 10.1002/humu.20040.

5. Dierks, T., Dickmanns, A., Preusser-Kunze, A., Schmidt, B., Mariappan, M., von Figura, K., Ficner, R., and Rudolph, M.G. (2005). Molecular basis for multiple sulfatase deficiency and mechanism for formylglycine generation of the human formylglycine-generating enzyme. Cell 121, 541–552. 10.1016/j.cell.2005.03.001.

6. Sardiello, M., Annunziata, I., Roma, G., and Ballabio, A. (2005). Sulfatases and sulfatase modifying factors: an exclusive and promiscuous relationship. Hum Mol Genet 14, 3203–3217. 10.1093/hmg/ddi351.

7. Schlotawa, L., Steinfeld, R., von Figura, K., Dierks, T., and Gartner, J. (2008). Molecular analysis of SUMF1 mutations: stability and residual activity of mutant formylglycine-generating enzyme determine disease severity in multiple sulfatase deficiency. Hum Mutat 29, 205. 10.1002/humu.9515.

8. Dierks, T., Schlotawa, L., Frese, M.A., Radhakrishnan, K., von Figura, K., and Schmidt, B. (2009). Molecular basis of multiple sulfatase deficiency, mucolipidosis II/III and Niemann-Pick C1 disease - Lysosomal storage disorders caused by defects of non-lysosomal proteins. Biochim Biophys Acta 1793, 710–725. 10.1016/j.bbamcr.2008.11.015.

9. Schlotawa, L., Ennemann, E.C., Radhakrishnan, K., Schmidt, B., Chakrapani, A., Christen, H.J., Moser, H., Steinmann, B., Dierks, T., and Gärtner, J. (2011). SUMF1 mutations affecting stability and activity of formylglycine generating enzyme predict clinical outcome in multiple sulfatase deficiency. Eur J Hum Genet 19, 253–261. 10.1038/ejhg.2010.219.

10. Schlotawa, L., Adang, L.A., Radhakrishnan, K., and Ahrens-Nicklas, R.C. (2020). Multiple Sulfatase Deficiency: A Disease Comprising Mucopolysaccharidosis, Sphingolipidosis, and More Caused by a Defect in Posttranslational Modification. Int J Mol Sci 21. 10.3390/ijms21103448.

11. Schlotawa, L., Dierks, T., Christoph, S., Cloppenburg, E., Ohlenbusch, A., Korenke, G.C., and Gärtner, J. (2019). Severe neonatal multiple sulfatase deficiency presenting with hydrops fetalis in a preterm birth patient. JIMD Rep 49, 48–52. 10.1002/jmd2.12074.

12. Ahrens-Nicklas, R., Schlotawa, L., Ballabio, A., Brunetti-Pierri, N., De Castro, M., Dierks, T., Eichler, F., Ficicioglu, C., Finglas, A., Gaertner, J., et al. (2018). Complex care of individuals with multiple sulfatase deficiency: Clinical cases and consensus statement. Mol Genet Metab 123, 337–346. 10.1016/j.ymgme.2018.01.005.

13. Schlotawa, L., Adang, L., De Castro, M., and Ahrens-Nicklas, R. (1993). Multiple Sulfatase Deficiency. In GeneReviews((R)), M.P. Adam, H.H. Ardinger, R.A. Pagon, S.E. Wallace, L.J.H. Bean, G. Mirzaa, and A. Amemiya, eds.

14. Adang, L.A., Schlotawa, L., Groeschel, S., Kehrer, C., Harzer, K., Staretz-Chacham, O., Silva, T.O., Schwartz, I.V.D., Gärtner, J., De Castro, M., et al. (2020). Natural history of multiple sulfatase deficiency: Retrospective phenotyping and functional variant analysis to characterize an ultra-rare disease. J Inherit Metab Dis 43, 1298–1309. 10.1002/jimd.12298.

15. Rigante, D., Cipolla, C., Basile, U., Gulli, F., and Savastano, M.C. (2017). Overview of immune abnormalities in lysosomal storage disorders. Immunol Lett 188, 79–85. 10.1016/j.imlet.2017.07.004.

16. Schlotawa, L., Preiskorn, J., Ahrens-Nicklas, R., Schiller, S., Adang, L.A., Gartner, J., and Friede, T. (2020). A systematic review and meta-analysis of published cases reveals the natural disease history in multiple sulfatase deficiency. J Inherit Metab Dis 43, 1288–1297. 10.1002/jimd.12282.

17. Sands, M.S., and Davidson, B.L. (2006). Gene therapy for lysosomal storage diseases. Mol Ther 13, 839–849. 10.1016/j.ymthe.2006.01.006.

18. Boucher, A.A., Miller, W., Shanley, R., Ziegler, R., Lund, T., Raymond, G., and Orchard, P.J. (2015). Long-term outcomes after allogeneic hematopoietic stem cell transplantation for metachromatic leukodystrophy: the largest single-institution cohort report. Orphanet J Rare Dis 10, 94. 10.1186/s13023-015-0313-y.

19. Aldenhoven, M., Wynn, R.F., Orchard, P.J., O’Meara, A., Veys, P., Fischer, A., Valayannopoulos, V., Neven, B., Rovelli, A., Prasad, V.K., et al. (2015). Long-term outcome of Hurler syndrome patients after hematopoietic cell transplantation: an international multicenter study. Blood 125, 2164–2172. 10.1182/blood-2014-11-608075.

20. Beschle, J., Doring, M., Kehrer, C., Raabe, C., Bayha, U., Strolin, M., Bohringer, J., Bevot, A., Kaiser, N., Bender, B., et al. (2020). Early clinical course after hematopoietic stem cell transplantation in children with juvenile metachromatic leukodystrophy. Mol Cell Pediatr 7, 12. 10.1186/s40348-020-00103-7.

21. Guffon, N., Pettazzoni, M., Pangaud, N., Garin, C., Lina-Granade, G., Plault, C., Mottolese, C., Froissart, R., and Fouilhoux, A. (2021). Long term disease burden post-transplantation: three decades of observations in 25 Hurler patients successfully treated with hematopoietic stem cell transplantation (HSCT). Orphanet J Rare Dis 16, 60. 10.1186/s13023-020-01644-w.

22. Fumagalli, F., Calbi, V., Natali Sora, M.G., Sessa, M., Baldoli, C., Rancoita, P.M.V., Ciotti, F., Sarzana, M., Fraschini, M., Zambon, A.A., et al. (2022). Lentiviral haematopoietic stem-cell gene therapy for early-onset metachromatic leukodystrophy: long-term results from a non-randomised, open-label, phase 1/2 trial and expanded access. The Lancet 399, 372–383. 10.1016/S0140-6736(21)02017-1.

23. Tan, E.Y., Boelens, J.J., Jones, S.A., and Wynn, R.F. (2019). Hematopoietic Stem Cell Transplantation in Inborn Errors of Metabolism. Front Pediatr 7, 433. 10.3389/fped.2019.00433.

24. Biffi, A., De Palma, M., Quattrini, A., Del Carro, U., Amadio, S., Visigalli, I., Sessa, M., Fasano, S., Brambilla, R., Marchesini, S., et al. (2004). Correction of metachromatic leukodystrophy in the mouse model by transplantation of genetically modified hematopoietic stem cells. J Clin Invest 113, 1118–1129. 10.1172/jci19205.

25. Biffi, A., Montini, E., Lorioli, L., Cesani, M., Fumagalli, F., Plati, T., Baldoli, C., Martino, S., Calabria, A., Canale, S., et al. (2013). Lentiviral hematopoietic stem cell gene therapy benefits metachromatic leukodystrophy. Science 341, 1233158. 10.1126/science.1233158.

26. Sergijenko, A., Langford-Smith, A., Liao, A.Y., Pickford, C.E., McDermott, J., Nowinski, G., Langford-Smith, K.J., Merry, C.L., Jones, S.A., Wraith, J.E., et al. (2013). Myeloid/Microglial driven autologous hematopoietic stem cell gene therapy corrects a neuronopathic lysosomal disease. Mol Ther 21, 1938–1949. 10.1038/mt.2013.141.

27. Ellison, S.M., Liao, A., Wood, S., Taylor, J., Youshani, A.S., Rowlston, S., Parker, H., Armant, M., Biffi, A., Chan, L., et al. (2019). Pre-clinical Safety and Efficacy of Lentiviral Vector-Mediated Ex Vivo Stem Cell Gene Therapy for the Treatment of Mucopolysaccharidosis IIIA. Mol Ther Methods Clin Dev 13, 399–413. 10.1016/j.omtm.2019.04.001.

28. Wada, M., Shimada, Y., Iizuka, S., Ishii, N., Hiraki, H., Tachibana, T., Maeda, K., Saito, M., Arakawa, S., Ishimoto, T., et al. (2020). Ex Vivo Gene Therapy Treats Bone Complications of Mucopolysaccharidosis Type II Mouse Models through Bone Remodeling Reactivation. Mol Ther Methods Clin Dev 19, 261–274. 10.1016/j.omtm.2020.09.012.

29. Miwa, S., Watabe, A.M., Shimada, Y., Higuchi, T., Kobayashi, H., Fukuda, T., Kato, F., Ida, H., and Ohashi, T. (2020). Efficient engraftment of genetically modified cells is necessary to ameliorate central nervous system involvement of murine model of mucopolysaccharidosis type II by hematopoietic stem cell targeted gene therapy. Mol Genet Metab 130, 262–273. 10.1016/j.ymgme.2020.06.007.

30. Gentner, B., Tucci, F., Galimberti, S., Fumagalli, F., De Pellegrin, M., Silvani, P., Camesasca, C., Pontesilli, S., Darin, S., Ciotti, F., et al. (2021). Hematopoietic Stem- and Progenitor-Cell Gene Therapy for Hurler Syndrome. N Engl J Med 385, 1929–1940. 10.1056/NEJMoa2106596.

31. Wakabayashi-Ito, N., and Nagata, S. (1994). Characterization of the regulatory elements in the promoter of the human elongation factor-1 alpha gene. J Biol Chem 269, 29831–29837.

32. Rivella, S., Callegari, J.A., May, C., Tan, C.W., and Sadelain, M. (2000). The cHS4 insulator increases the probability of retroviral expression at random chromosomal integration sites. J Virol 74, 4679–4687. 10.1128/jvi.74.10.4679-4687.2000.

33. Zufferey, R., Donello, J.E., Trono, D., and Hope, T.J. (1999). Woodchuck hepatitis virus posttranscriptional regulatory element enhances expression of transgenes delivered by retroviral vectors. J Virol 73, 2886–2892. 10.1128/jvi.73.4.2886-2892.1999.

34. Breda, L., Ghiaccio, V., Tanaka, N., Jarocha, D., Ikawa, Y., Abdulmalik, O., Dong, A., Casu, C., Raabe, T.D., Shan, X., et al. (2021). Lentiviral vector ALS20 yields high hemoglobin levels with low genomic integrations for treatment of beta-globinopathies. Mol Ther 29, 1625–1638. 10.1016/j.ymthe.2020.12.036.

35. Schlotawa, L., Tyka, K., Kettwig, M., Ahrens-Nicklas, R.C., Baud, M., Berulava, T., Brunetti-Pierri, N., Gagne, A., Herbst, Z.M., Maguire, J.A., et al. (2023). Drug screening identifies tazarotene and bexarotene as therapeutic agents in multiple sulfatase deficiency. EMBO Mol Med 15, e14837. 10.15252/emmm.202114837.

36. Herbst, Z.M., Hong, X., Urdaneta, L., Klein, T., Waggoner, C., Liao, H.C., Kubaski, F., Giugliani, R., Fuller, M., and Gelb, M.H. (2023). Endogenous, non-reducing end glycosaminoglycan biomarkers are superior to internal disaccharide glycosaminoglycan biomarkers for newborn screening of mucopolysaccharidoses and GM1 gangliosidosis. Mol Genet Metab, 107632. 10.1016/j.ymgme.2023.107632.

37. Saville, J.T., McDermott, B.K., Fletcher, J.M., and Fuller, M. (2019). Disease and subtype specific signatures enable precise diagnosis of the mucopolysaccharidoses. Genet Med 21, 753–757. 10.1038/s41436-018-0136-z.

38. Sorrentino, N.C., Presa, M., Attanasio, S., Cacace, V., Sofia, M., Zuberi, A., Ryan, J., Ray, S., Petkovic, I., Radhakrishnan, K., et al. (2023). New mouse models with hypomorphic SUMF1 variants mimic attenuated forms of multiple sulfatase deficiency. J Inherit Metab Dis 46, 335–347. 10.1002/jimd.12577.

39. Settembre, C., Annunziata, I., Spampanato, C., Zarcone, D., Cobellis, G., Nusco, E., Zito, E., Tacchetti, C., Cosma, M.P., and Ballabio, A. (2007). Systemic inflammation and neurodegeneration in a mouse model of multiple sulfatase deficiency. Proc Natl Acad Sci U S A 104, 4506–4511. 10.1073/pnas.0700382104.

40. Guyenet, S.J., Furrer, S.A., Damian, V.M., Baughan, T.D., La Spada, A.R., and Garden, G.A. (2010). A simple composite phenotype scoring system for evaluating mouse models of cerebellar ataxia. J Vis Exp. 10.3791/1787.

41. Hong, X., Daiker, J., Sadilek, M., Ruiz-Schultz, N., Kumar, A.B., Norcross, S., Dansithong, W., Suhr, T., Escolar, M.L., Ronald Scott, C., et al. (2021). Toward newborn screening of metachromatic leukodystrophy: results from analysis of over 27,000 newborn dried blood spots. Genet Med 23, 555–561. 10.1038/s41436-020-01017-5.

42. Pitts, M.W. (2018). Barnes Maze Procedure for Spatial Learning and Memory in Mice. Bio Protoc 8. 10.21769/bioprotoc.2744.

43. Lalonde, R., and Strazielle, C. (2011). Brain regions and genes affecting limb-clasping responses. Brain Res Rev 67, 252–259. 10.1016/j.brainresrev.2011.02.005.

44. Herbst, Z.M., Urdaneta, L., Klein, T., Burton, B.K., Basheeruddin, K., Liao, H.C., Fuller, M., and Gelb, M.H. (2022). Evaluation of Two Methods for Quantification of Glycosaminoglycan Biomarkers in Newborn Dried Blood Spots from Patients with Severe and Attenuated Mucopolysaccharidosis Type II. Int J Neonatal Screen 8. 10.3390/ijns8010009.

45. Adang, L.A., Mowafy, S., Herbst, Z.M., Zhou, Z., Schlotawa, L., Radhakrishnan, K., Bentley, B., Pham, V., Yu, E., Pillai, N.R., et al. (2023). Biochemical signatures of disease severity in multiple sulfatase deficiency. J Inherit Metab Dis. 10.1002/jimd.12688.

46. Tucci, F., Consiglieri, G., Cossutta, M., and Bernardo, M.E. (2023). Current and Future Perspective in Hematopoietic Stem Progenitor Cell-gene Therapy for Inborn Errors of Metabolism. Hemasphere 7, e953. 10.1097/hs9.0000000000000953.

47. Breda, L., Casu, C., Gardenghi, S., Bianchi, N., Cartegni, L., Narla, M., Yazdanbakhsh, K., Musso, M., Manwani, D., Little, J., et al. (2012). Therapeutic hemoglobin levels after gene transfer in β-thalassemia mice and in hematopoietic cells of β-thalassemia and sickle cells disease patients. PLoS One 7, e32345. 10.1371/journal.pone.0032345.

48. Goodman, M.A., Arumugam, P., Pillis, D.M., Loberg, A., Nasimuzzaman, M., Lynn, D., van der Loo, J.C.M., Dexheimer, P.J., Keddache, M., Bauer, T.R., Jr., et al. (2018). Foamy Virus Vector Carries a Strong Insulator in Its Long Terminal Repeat Which Reduces Its Genotoxic Potential. J Virol 92. 10.1128/jvi.01639-17.

49. Liu, M., Maurano, M.T., Wang, H., Qi, H., Song, C.Z., Navas, P.A., Emery, D.W., Stamatoyannopoulos, J.A., and Stamatoyannopoulos, G. (2015). Genomic discovery of potent chromatin insulators for human gene therapy. Nat Biotechnol 33, 198–203. 10.1038/nbt.3062.

50. Tucci, F., Galimberti, S., Naldini, L., Valsecchi, M.G., and Aiuti, A. (2022). A systematic review and meta-analysis of gene therapy with hematopoietic stem and progenitor cells for monogenic disorders. Nat Commun 13, 1315. 10.1038/s41467-022-28762-2.

51. Pillai, N.R., Orchard, P.J., Ahrens-Nicklas, R., Adang, L., Elsea, S.H., and Whitley, C.B. (2021). Evaluation of the effectiveness of hematopoietic stem cell transplantation in multiple sulfatase deficiency. Molecular Genetics and Metabolism 132, S87. 10.1016/j.ymgme.2020.12.207.

52. Presa, M., Ryan, J., and Lutz, C. (2023). Efficacy of syngeneic bone marrow transplant for the treatment of multiple sulfatase deficiency. Molecular Genetics and Metabolism 138, 107278.

53. Spampanato, C., De Leonibus, E., Dama, P., Gargiulo, A., Fraldi, A., Sorrentino, N.C., Russo, F., Nusco, E., Auricchio, A., Surace, E.M., and Ballabio, A. (2011). Efficacy of a combined intracerebral and systemic gene delivery approach for the treatment of a severe lysosomal storage disorder. Mol Ther 19, 860–869. 10.1038/mt.2010.299.

54. Bailey, R., Presa, M., Ray, S., Bailey, L., Coombs, H., Walls, R., Lutz, C., and Gray, S. (2021). Preclinical studies to support the intrathecal delivery of scAAV9/SUMF1 as a gene replacement therapy for multiple sulfatase deficiency. Molecular Genetics and Metabolism 132, S17–S18. 10.1016/j.ymgme.2020.12.020.

55. Baum, H., Dodgson, K.S., and Spencer, B. (1959). The assay of arylsulphatases A and B in human urine. Clin Chim Acta 4, 453–455. 10.1016/0009-8981(59)90119-6.

56. Karpova, E.A., Voznyi Ya, V., Keulemans, J.L., Hoogeveen, A.T., Winchester, B., Tsvetkova, I.V., and van Diggelen, O.P. (1996). A fluorimetric enzyme assay for the diagnosis of Sanfilippo disease type A (MPS IIIA). J Inherit Metab Dis 19, 278–285. 10.1007/bf01799255.

