## Supplemental Material for "Hematopoietic stem cell gene therapy improves outcomes in a clinically relevant mouse model of Multiple Sulfatase Deficiency"

**Figure S1. *SUMF1 ex vivo* gene therapy does not rescue astrocytosis in the cortex.**

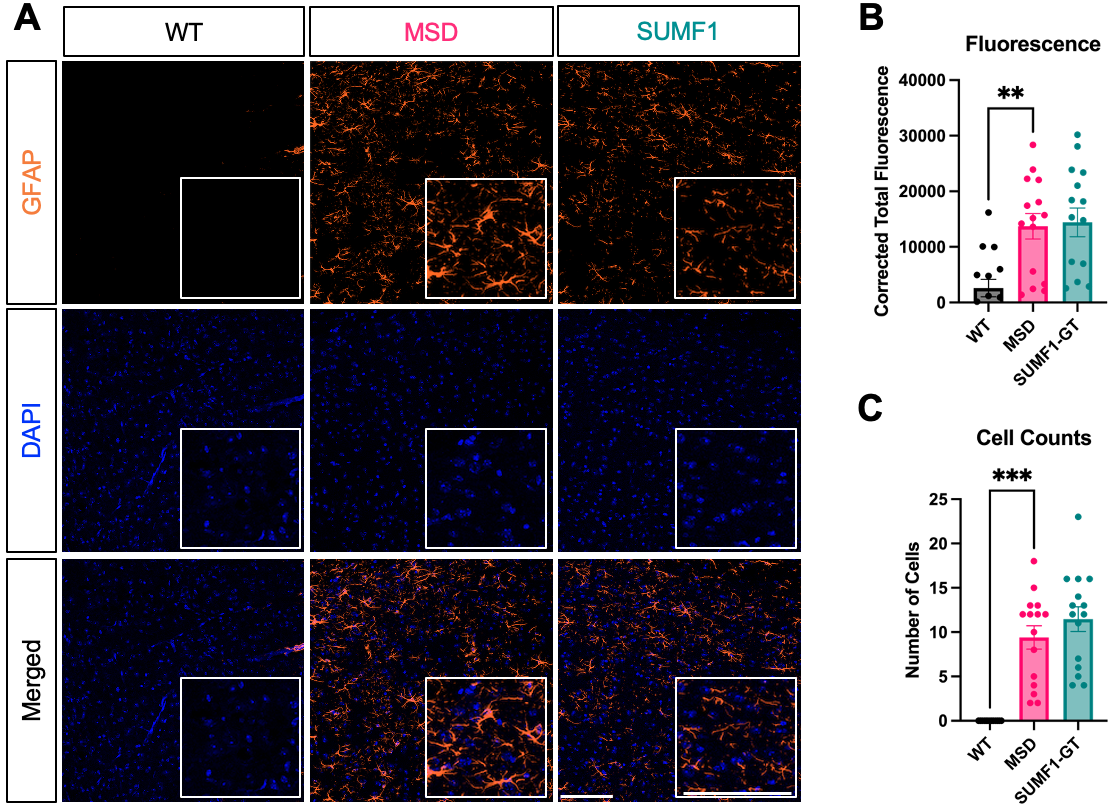

A) Representative immunofluorescence images of GFAP+ cells in cortical brain sections of WT mice (WT), untreated MSD mice (MSD), and MSD mice receiving *SUMF1* HSCT-GT (SUMF1-GT) reveal greater astrocytosis in untreated and treated MSD mice compared to WT. Labeling with anti-GFAP antibody (orange pseudo-fluorescence) and DAPI (nuclei, blue). Scale bars = 100 μm. B) Quantification of GFAP immunofluorescence intensity confirms significant increase in untreated and treated MSD animals. C) Quantification of GFAP+ cells in cortical sections validates increased astrocytosis in untreated and treated MSD mice. For all quantifications, three biological replicates were analyzed with 5 images per replicate, one-way ANOVA, followed by Bonferroni’s multiple comparisons test to compare each group to the untreated MSD control group. For all panels: ** *p* < 0.01, *** *p* <0.001. Data are represented as mean ± SEM.

**Figure S2. MSD mice do not exhibit deficits in open field or elevated zero tests.**

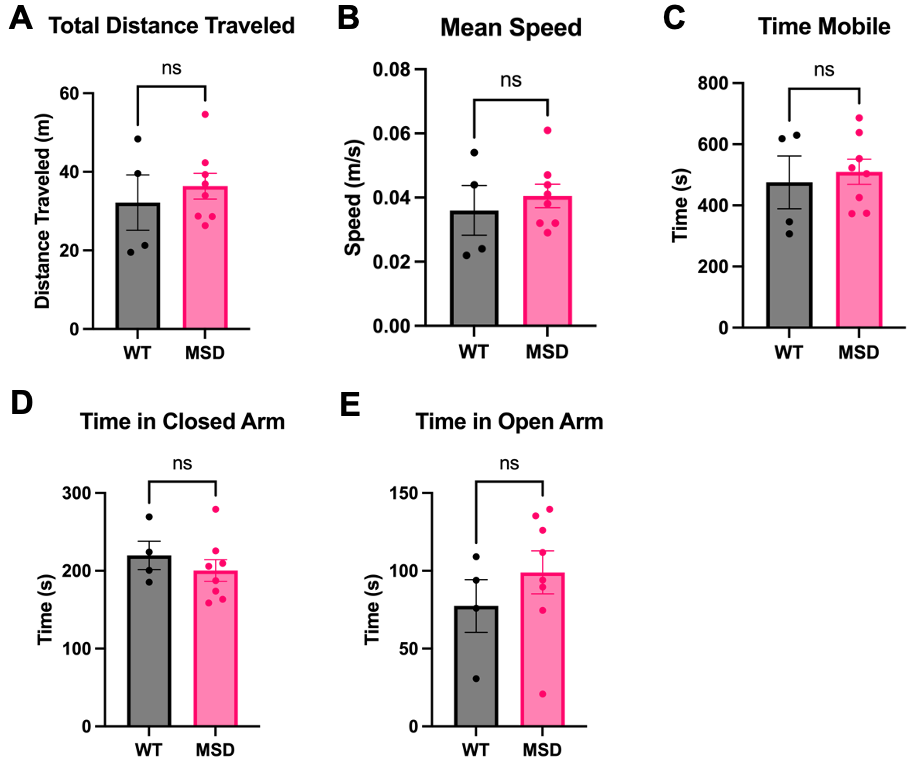

A) Quantification of the total distance traveled (meters) in the open field test for WT and MSD mice. B) Quantification of the mean speed (meters/second) in the open field test for WT and MSD mice. C) Quantification of the time mobile (seconds) in the open field test for WT and MSD mice. D) Quantification of the time in closed arm (seconds) in the elevated zero test for WT and MSD mice. E) Quantification of the time in open arm (seconds) in the elevated zero test for WT and MSD mice. For all panels, *N* = 4-8 mice, unpaired t-test. Data are represented as mean ± SEM.
